# Atomistic basis of microtubule dynamic instability assessed via multiscale modeling

**DOI:** 10.1101/2020.01.07.897439

**Authors:** Mahya Hemmat, David J. Odde

## Abstract

Microtubule “dynamic instability,” the abrupt switching from assembly to disassembly caused by the hydrolysis of GTP to GDP within the β subunit of the αβ-tubulin heterodimer, is necessary for vital cellular processes such as mitosis and migration. Despite existing high-resolution structural data, the key mechanochemical differences between the GTP and GDP states that mediate dynamic instability behavior remain unclear. Starting with a published atomic-level structure as an input, we used multiscale modeling to find that GTP hydrolysis results in both longitudinal bond weakening (~4 k_B_T) and an outward bending preference (~1.5 k_B_T) to both drive dynamic instability and give rise to the microtubule tip structures previously observed by light and electron microscopy. More generally, our study provides an example where atomic level structural information is used as the sole input to predict cellular level dynamics without parameter adjustment.

## Introduction

Biological processes occur over a wide range of spatial and temporal scales, spanning from angstroms and picoseconds at the atomistic level to micrometers and minutes at the cellular level. While rapid advances in structural biology are yielding unprecedented insight into biological structures at the angstrom scale, converting this information into quantitative prediction of cellular level dynamics on the time scale of minutes remains a major challenge. Fortunately, several multiscale modeling approaches have been developed to address this challenge (1–8). For example, association kinetics of small biochemical ligand-receptor systems have been explored computationally by a multiscale approach that matches experimental values, connecting molecular dynamics (MD) simulations to Brownian dynamics (BD) simulations with milestoning theory (9). Another example of a multi-scale framework initiating from atomistic details studied actin filaments by integrating all-atom MD simulations and coarse-grained (CG) techniques to study the impact of the nucleotide state on the filament’s conformation, and the hydrolysis rate constant (2,10). These methods, although bridging information across two or three scales temporally and spatially, do not yet provide a single framework to connect across length-time scales from atoms to cells.

Microtubules, are a prime example of a complex biological system with spatial and temporal scales ranging from atomistic phenomena such as nucleotide hydrolysis to cellular level organization and behavior that mediate cellular functions such as cell division and migration (11). Microtubules self-assemble from αβ-tubulin heterodimers that form both longitudinal and lateral bonds to build a hollow cylinder (12–14). Tubulin heterodimers have conserved protein structures with defined binding zones which makes them ideal to study at the atomistic-molecular level (15–17). A key feature of microtubule assembly is the phenomenon of “dynamic instability,” where the highly dynamic plus-end switches abruptly and stochastically between extended phases of net growth and net shortening, which depends on the guanosine triphosphate/diphosphate (GTP/GDP) nucleotide state in the β-tubulin subunit (18,19). Whereas microtubules grow via net addition of GTP-tubulin subunits, the GTP-tubulin soon hydrolyzes within the microtubule lattice to GDP-tubulin resulting in a so-called “GTP cap” that stabilizes the growing microtubule (20–23). When the GTP cap is lost through a combination of GTP hydrolysis and stochastic loss of GTP subunits, the labile GDP-tubulin core of the microtubule is exposed and the microtubule undergoes “catastrophe” followed by rapid shortening (12). Because dynamic instability is essential for mitosis and cell migration, a key question is what are the fundamental thermodynamic and mechanical differences between the GTP and GDP states of the tubulin subunit that lead to net growth and shortening, respectively.

Previous studies of microtubule dynamic instability posited that the energetic difference between the two nucleotide states of tubulin in models of microtubule assembly (24,25,34–37,26–33) is due to differences in the tubulin-tubulin lateral bond strength (25,27,28,31,32,34), tubulin-tubulin longitudinal bond strength (31,32,37), the bending preference or flexibility (24,26,29,30,33,35,36), or indirectly dependent on the lateral bond as a result of nucleotide-dependent bending strain energy (38). At the level of atomic structure, a recent cryo-electron microscopy (cryo-EM) study showed that GDP, GDP-Pi and GTPγS all have compacted lattices (~82 Å) compared to the GMPCPP extended lattice (~ 83.7 Å) (32), consistent with conclusions from earlier cryo-EM studies (37,39). This study also suggested that the compaction in GDP-lattice causes stronger longitudinal interactions and perturbs the lateral bond between the protofilaments, thus resulting in splayed protofilament ends and microtubule depolymerization. However, in this, and the other structural studies cited above, the conclusions drawn from cryo-EM studies regarding the mechanism of dynamic instability are based on the assumption that the contacts of the residues remain stationary as captured by the cryo-EM structure.

To gain insight into the atomistic dynamics, MD simulations have been performed using the protein data bank (PDB) structures as the sole input. Using the structure of tubulin protofilaments as a function of nucleotide state, a multiscale approach was taken to integrate the results of 100 ns equilibrium MD simulation with a CG-MD analysis (31), representing a molecular group of atoms as one particle (1). This study concluded that GDP-microtubules are less stable because GDP-tubulin has weaker longitudinal and lateral contacts in the lattice compared to GTP-tubulin. However, this conclusion was drawn from MD analysis of contact maps and CG analysis of bond strength and lengths, while more recent work (40) using 10 all-atom MD replicates per nucleotide state combined with BD and thermokinetic modeling revealed no evidence of nucleotide dependence of the lateral bond. More generally, as stated by the authors (31), analysis of residue contact map and CG bond strength and length is not a measure of the actual binding strength of tubulin dimers; rather, an energy landscape of the lateral and longitudinal interaction of the dimers needs to be calculated. In addition, while a number of other MD studies have contributed toward an atomistic understanding of *intradimer* bending mechanics (29,30), to our knowledge, previous studies have not examined the free energy of *interdimer* bending as a function of its nucleotide besides equilibrium trajectory analysis (35,36,41). Finally, previous studies have not estimated the nucleotide dependence of the strength of the longitudinal bond, generally regarded as stronger than the lateral bond. Thus, an integrated multiscale dynamics approach is needed to understand the atomistic basis of microtubule dynamic instability, and, more generally, to connect atomic structures to cellular level behavior.

To address this challenge, we performed simulations at multiple scales where the output of one scale was used as the input to the scale above it. Using published crystal structures as inputs, we performed MD simulations of tubulin-tubulin interactions to obtain an estimated longitudinal potential of mean force (PMF), with which we then simulated tubulin addition and loss from the microtubule lattice via BD to obtain estimated k_on_, k_off_, and ΔG^0^. Using these k_on_ and k_off_ values as inputs, we used thermokinetic and mechanochemical modeling to predict microtubule dynamic instability at the scale of micrometers and minutes. Thus, we were able to simulate the cellular dynamics using the published crystal structures as the only inputs with no adjustable parameters. We find that the GTP-tubulin longitudinal bond has a stronger PMF than GDP-tubulin by ΔU_long_≈ 6.6 ± 2.8 k_B_T, which translates into a standard Gibbs free energy difference of ≈ 4 ±0.5 k_B_T. While this difference largely explains the atomistic basis of dynamic instability, we further found via thermokinetic and mechanochemical modeling that an outward bending preference of GDP-tubulin relative to GTP-tubulin contributes to dynamic instability and is necessary for realistic tip structures (~1.5 k_B_T). Overall, our multi-scale approach (Fig 1) presents a methodology, using microtubule dynamic instability as a specific example, for using atomistic structural information to predict cellular level behaviors (42) without parameter adjustment.

**Figure 1.**
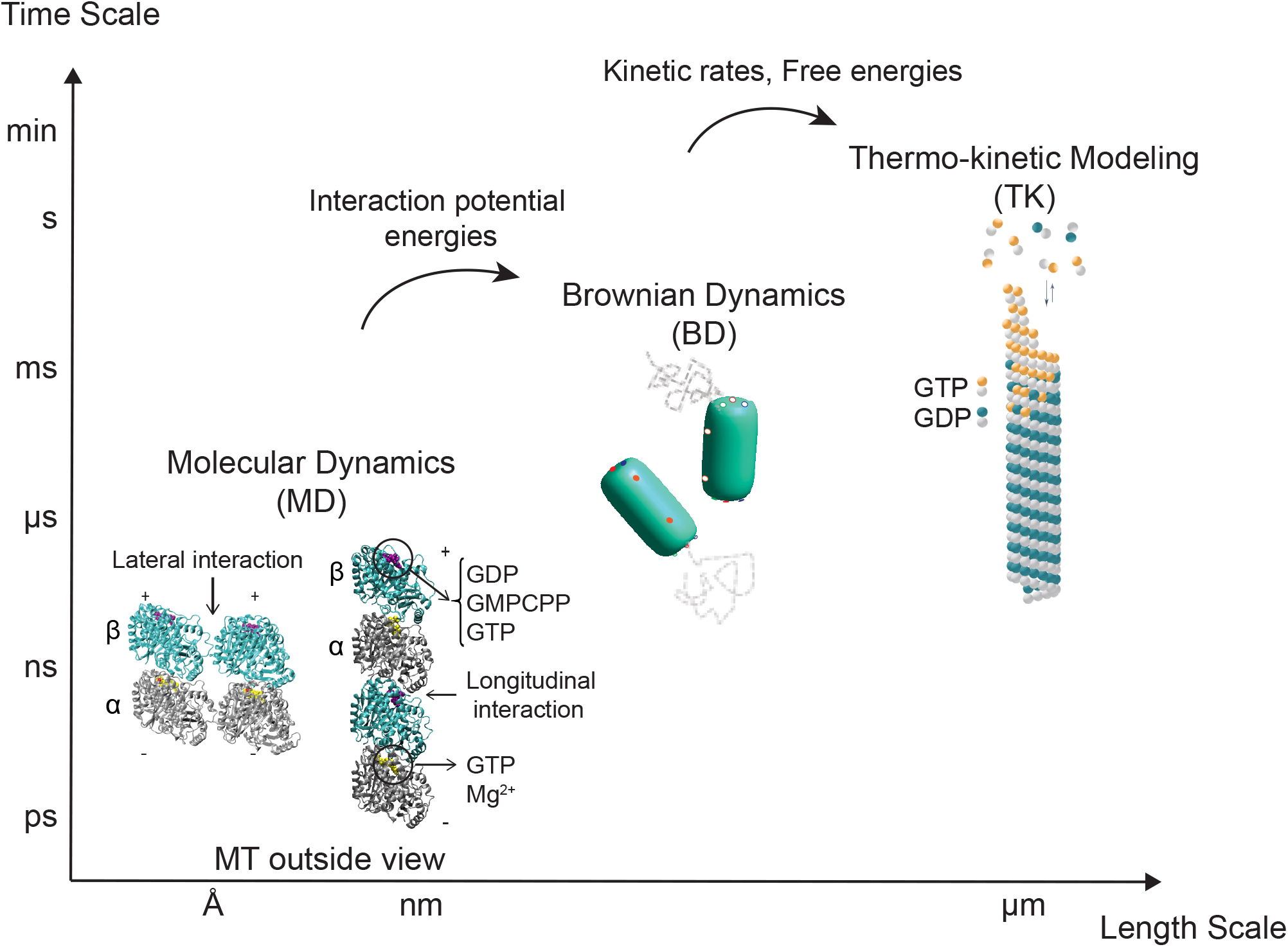
Multi-scale modeling approach to study microtubule dynamics from atoms, Å/ps length/time scales, to microtubules, µm/min length/time scales.

## Results

### Tubulin dimers reach equilibrium in the absence and presence of microtubule lattice constraints

To investigate the longitudinal interaction between tubulin heterodimers, an interaction essential to protofilament formation, we modeled a pair of tubulin dimers stacked longitudinally via MD simulations. Since tubulin structures were initially obtained from straight protofilaments (37) (Fig 2A), we were interested to see how removing the lattice constraints would influence oligomer (dimer of heterodimers) bending and conformation. During the simulation, we monitored the root-mean-square deviation (RMSD) of the backbone atoms to ensure global equilibrium (Fig 2B). We also considered the possibility that microtubule lattice constraints affect the conformations of the dimers, mostly by confining their bending motions. These constraints are believed to be stored as strain energy that keeps the dimer’s conformation straight in the lattice. To test this hypothesis, we simulated the effects of lateral neighbors and the bottom longitudinal neighbor as harmonic constraints on the atoms mainly involved in the lateral (40) and longitudinal bond (highlighted in red in Fig 2C). As for harmonic stiffness, we first calculated the longitudinal PMF for unconstrained dimers and then used that along with our previously calculated lateral PMF (40) to estimate a stiffness for the bonds by fitting a harmonic potential around the potential well. We obtained stiffnesses of *κ*=1 and 0.6 kcal/mol/Å^2^ (*κ*=0.7 and 0.4 nN/nm) for lateral and longitudinal bonds, respectively. In this way, we simulated the extreme case of the lattice constraints for a dimer at a protofilament tip, which is having two lateral neighbors and one longitudinal neighbor.

**Figure 2.**
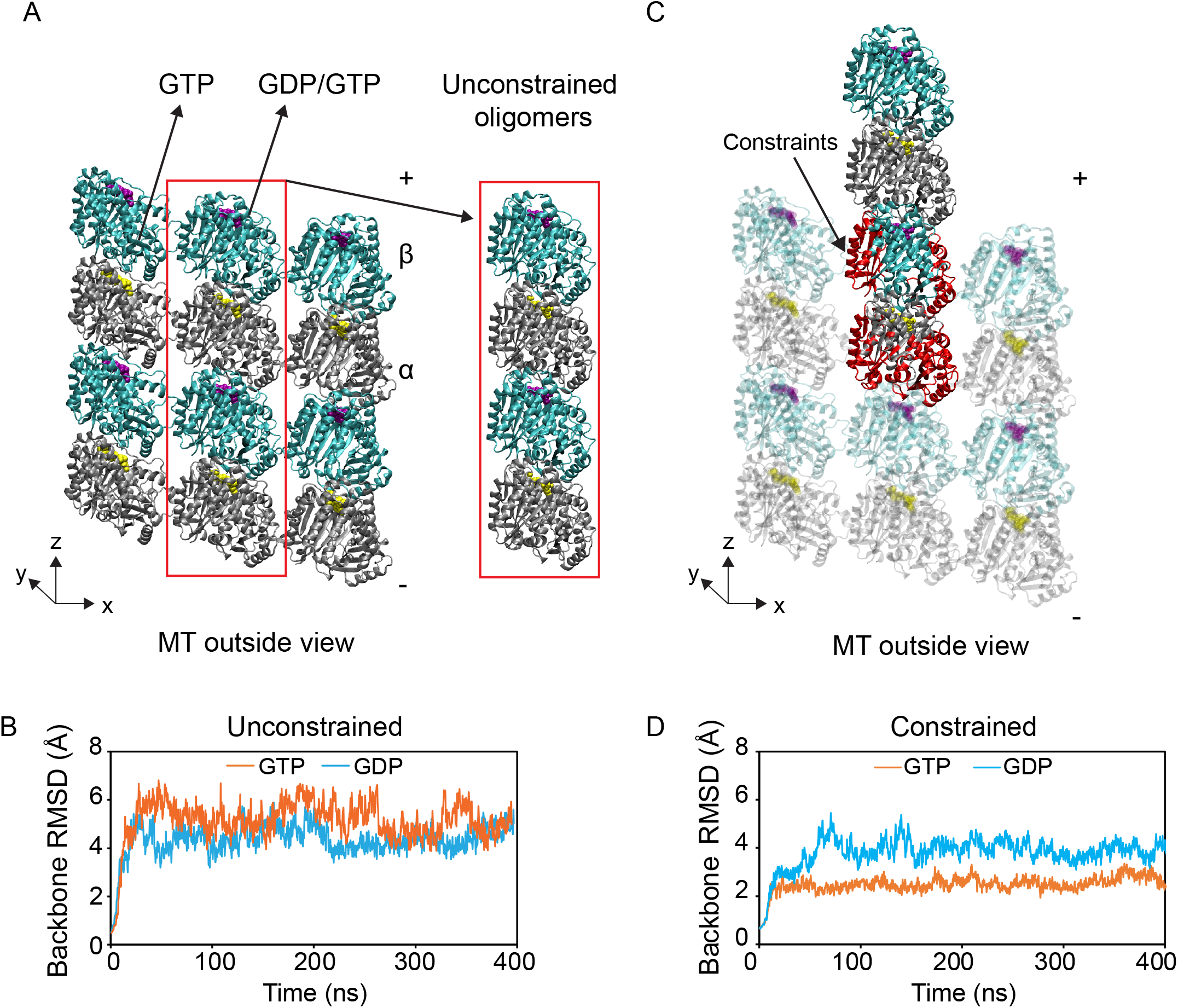
MD simulation systems for (A) unconstrained and (C) constrained longitudinal tubulin-tubulin interactions (PDB ID: 3JAS, 3JAT) with their backbone RMSD. Backbone RMSDs in angstroms for the 400 ns trajectories for both GDP and GTP nucleotide states show the equilibrium state for (B) unconstrained and (D) constrained simulations. Silver shows α-subunit, cyan shows GDP-β subunit and orange is GTP-β subunit. Red atoms show lattice constrained atoms.

We found that these lattice constraints do not deform the protein structure, as evidenced by the plateau of the RMSD of the backbone atoms after ~70ns for both nucleotide states (Fig 2D). Our simulations confirmed the two major bending modes for the oligomers, tangential and radial outward bending, regardless of the nucleotide or lattice constraints (Fig S1A, Supplemental Movies S1 and S2), as previously shown for single tubulin dimers (29,40). To compare our data to previously published tubulin MD simulations (29,35), the last 300ns of our trajectories were analyzed further in terms of root mean squared fluctuations (RMSF) and bending angle dynamics (Fig S1-S3, Table S1A, S1B, S2A, S2B), which confirmed behavior consistent with these previous studies. We avoid drawing strong quantitative conclusions about nucleotide dependence of bending angles based solely on these results since the angle data is highly variable and is not likely to be converged due to longer relaxation times. Nonetheless, we can conclude that our protein backbone structure is globally equilibrated, and the presence of the lattice constrains the fluctuations of the dimers, resulting in a straighter oligomer configuration, as evidenced by the reduction of the average backbone RMSD (Fig 2B, 2D) and average structures obtained from the trajectories (Fig S1B). The global equilibration allowed us to proceed to probe the non-covalent binding nature of the equilibrated longitudinal interface and the longitudinal bond strength, i.e. the longitudinal PMF.

### GTP- and GDP-tubulin have two longitudinal interaction zones in common but differ on a third zone

High-resolution cryo-EM microtubule structures identified three main longitudinal contact zones between the dimers in a protofilament (43). Three zones also reproduced the best estimates of experimental on-off rates of protein assembly when modeled via BD simulation (44,45). To investigate whether tubulin oligomers would maintain their longitudinal contacts despite the initial relaxation of the lattice compaction observed in GDP-lattice upon equilibration (Fig S4) (37), we searched for residues involved (>25%) in H-bond and ionic interactions at the interdimer interface. Our computational results, shown in Table 1 and Fig 3A and 3B, identified three non-symmetrical longitudinal interaction zones in each equilibrated nucleotide state in the case where the bottom dimer is lattice-constrained: 1) S9 and H10-S9 loop with T5 loop, 2) H11-H11’, H11’, H11’-H12 with H8-S7 and H4-S5 loops, and 3T) H6 with H10 in GTP-tubulin only and 3D) residues 2 to 4 in the α-subunit’s N-terminus with T2 loop in GDP-tubulin only. In contrast to the results of Nogales et al. (1999), the interaction of T7 loop with T2, T1, and the nucleotides (termed “zone C” by Nogales *et al.*) was less than 10% and 18% of the total trajectories in GDP and GTP states, respectively. This is due to the relaxation of the simulated dimers in water without the constraints of the lattice around all the dimers, as clearly observed in the compaction release of GDP dimers, which increases the distance between the nucleotide and interacting residues.

**Table 1.** Interdimer longitudinal zone interactions in constrained tubulin dimers

| Nucleotide state | Interaction | Secondary structure | % Occupancy* | Zone |
| --- | --- | --- | --- | --- |
| GDP | H-bond | N-terminus ARG2 --- T2 loop | 77.5% | 3D |
|  |  | H8 ---T3 loop | 55.8% | 2 |
|  |  | H8-S7 loop --- H11-H11' | 54.2% | 2 |
|  |  | H10-S9, S9 --- T5 loop* | 31.5 % | 1 |
|  | Salt bridge | H11'-H12 --- H4-S5* | 65.3% | 2 |
| GTP | H-bond | H8-S7 loop --- H11-H11', H11' | 38.6% | 2 |
|  |  | S9 --- T5 loop | 29.4 % | 1 |
|  | Salt bridge | H6 --- H10 | 42.8% | 3T |

\* < 25% in the unconstrained dimers

**Figure 3.**
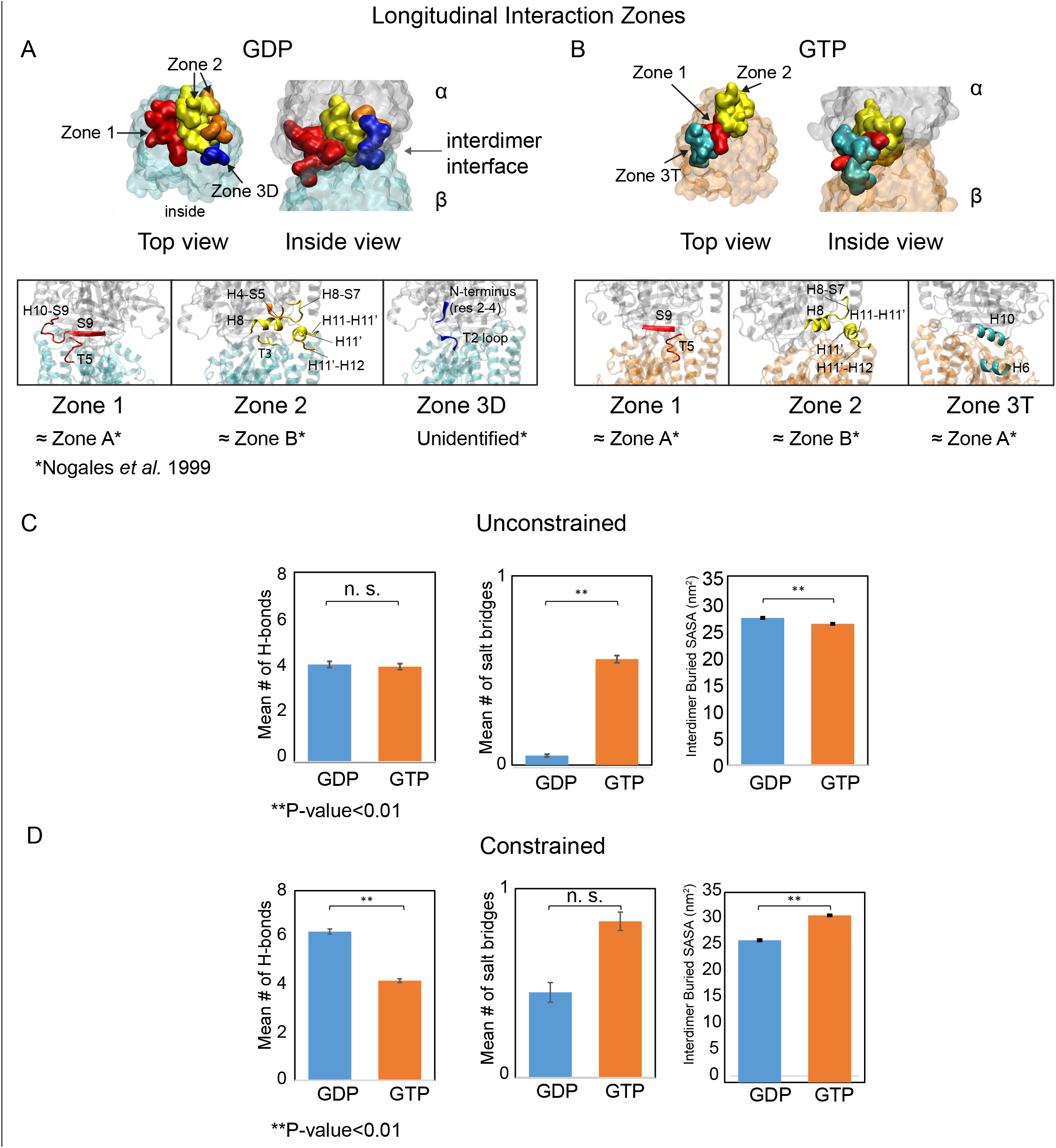
Three main longitudinal interaction zones are identified for GDP- and GTP-tubulins, and decomposed as H-bonds, salt bridges and hydrophobic interactions. (A, B) longitudinal zones are highlighted at the interdimer interface as red (zone 1), yellow and orange (zone 2), dark blue (zone 3D) and cyan (zone 3T) for GDP- and GTP-states, respectively. Panels show a more detailed view of the secondary structures involved in the interaction zones (* shows the equivalent zones identified by Nogales *et al.* 1999). (C) Mean number of H-bonds, salt bridges, and BSASA involved at the interdimer interface for more than 25% for GDP- and GTP-tubulins in unconstrained and (D) constrained simulations. N.S. is not significant (p >0.05).

These results also suggest that GTP- and GDP-dimers differ in one of their major longitudinal interaction zones, potentially causing different longitudinal bond strengths as a function of nucleotide state. To identify the relative contribution of various non-covalent interaction types at the longitudinal interface, we calculated the number of H-bonds, salt-bridges and extent of hydrophobic interactions quantified as buried solvent accessible surface area (BSASA) involved (>25%) in the interdimer interaction, as shown in Fig 3C and 3D. We found that the longitudinal interface is dominated by hydrophobic interactions (~25-30 nm^2^ BSASA), which is qualitatively distinct from the lateral interface which is dominated by H-bonds and ionic interactions (40). As expected, lattice constraints make the number of longitudinal contacts higher by preventing the dimers from bending considerably. However, in contrast to previous studies which argue that higher contact numbers equals a stronger bond (32), we note that this metric is not necessarily representative of the total longitudinal bond strength and is more an indicator of the quantity of longitudinal contacts without knowing their relative strengths. While these analyses inform our qualitative understanding of the nature of the bond and the relative shifts that occur upon nucleotide hydrolysis, assessing the quantitative strength of a bond requires calculating bond potentials (PMFs) and comparing the potential well-depths.

### The longitudinal bond is significantly stronger for GTP-tubulin than it is for GDP-tubulin

The differential strength of the longitudinal interactions between GTP- and GDP-tubulin heterodimers could contribute significantly to the phenomenon of dynamic instability. Therefore, we estimated the PMF of the longitudinal center of mass (COM) to COM distance for GDP- and GTP-tubulins, as free oligomers or lattice constrained oligomers. Our distance-based reaction coordinate choice, although limited to low dimer rotational degrees, was found to have the highest efficiency of longitudinal binding (Fig S5) and is more common in the literature (46). We used umbrella sampling with WHAM (47,48) (see Methods) to efficiently sample the ensemble, which has the advantage over traditional time-averaging techniques of overcoming energy barriers inaccessible to traditional MD simulations. To make conclusions about bond strengths on the scale of k_B_T, we found it necessary to run multiple replicates (Table S3). As shown in Fig 4A and 4B, our longitudinal PMFs clearly show a stronger GTP tubulin-tubulin interaction compared to GDP, regardless of the presence of lattice constraints. We found that the potential well-depths for the two nucleotide states differ by ~6.6 *k*_B_T (p<0.02; Table 2).

**Table 2.**
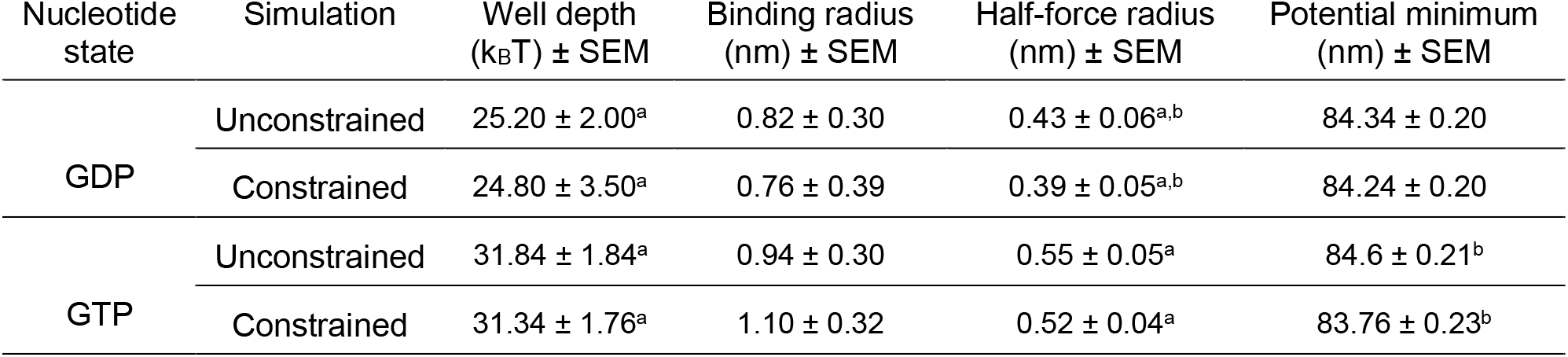
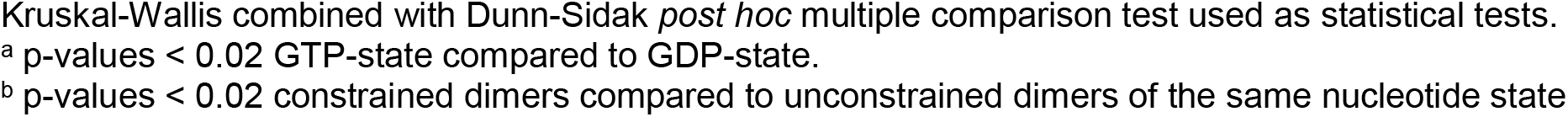
Potential interaction parameter values ± SEM of 10 replicates for unconstrained tubulin dimers and 5 replicates for constrained tubulin dimers

| Nucleotide state | Simulation | Well depth ($k_B T$) ± SEM | Binding radius (nm) ± SEM | Half-force radius (nm) ± SEM | Potential minimum (nm) ± SEM |
| --- | --- | --- | --- | --- | --- |
| GDP | Unconstrained | 25.20 ± 2.00<sup>a</sup> | 0.82 ± 0.30 | 0.43 ± 0.06<sup>a,b</sup> | 84.34 ± 0.20 |
| | Constrained | 24.80 ± 3.50<sup>a</sup> | 0.76 ± 0.39 | 0.39 ± 0.05<sup>a,b</sup> | 84.24 ± 0.20 |
| GTP | Unconstrained | 31.84 ± 1.84<sup>a</sup> | 0.94 ± 0.30 | 0.55 ± 0.05<sup>a</sup> | 84.6 ± 0.21<sup>b</sup> |
| | Constrained | 31.34 ± 1.76<sup>a</sup> | 1.10 ± 0.32 | 0.52 ± 0.04<sup>a</sup> | 83.76 ± 0.23<sup>b</sup> |

Kruskal-Wallis combined with Dunn-Sidak *post hoc* multiple comparison test used as statistical tests.
<sup>a</sup> p-values < 0.02 GTP-state compared to GDP-state.
<sup>b</sup> p-values < 0.02 constrained dimers compared to unconstrained dimers of the same nucleotide state

**Figure 4.**
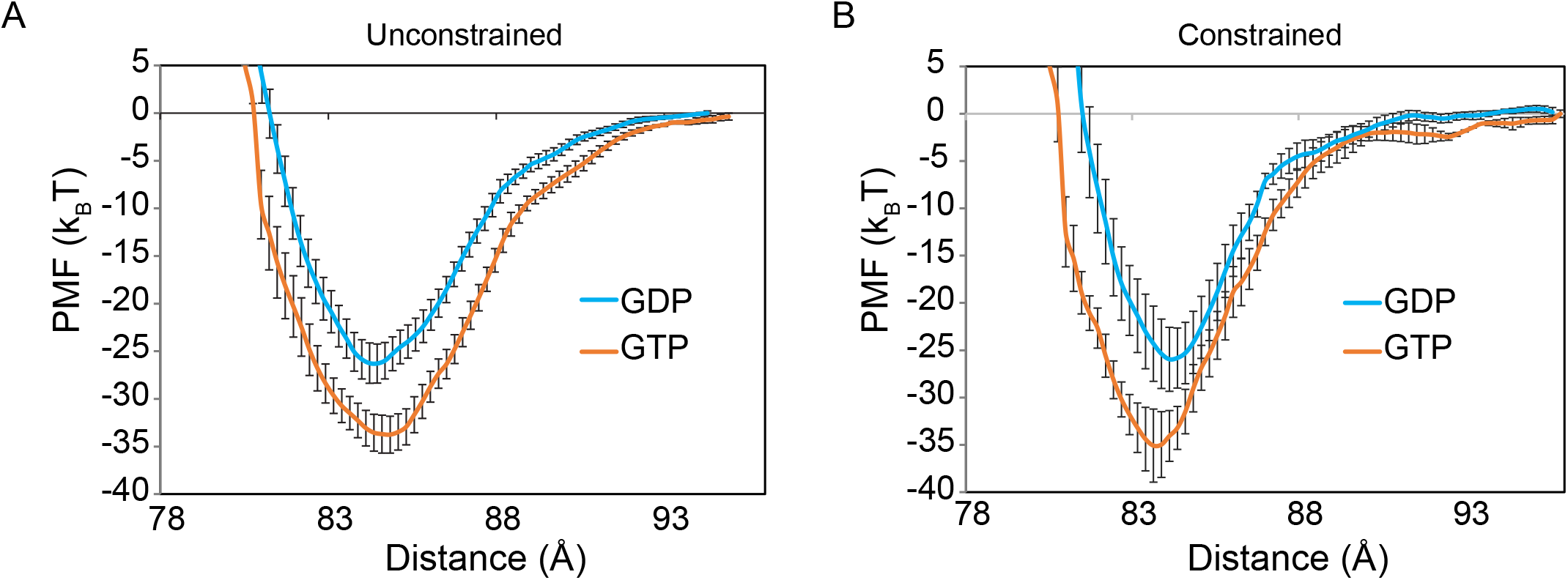
GTP-tubulin has a stronger longitudinal interaction potential compared to GDP-tubulin in both constrained and unconstrained simulations. (A) Average longitudinal PMFs as a function of COM-to-COM distance for both nucleotide states for unconstrained, and (B) constrained conditions. Error bars show standard error of the mean of 10 replicates for unconstrained and 5 replicates for constrained simulations.

To investigate the stiffnesses of the potentials, we calculated the half-force radius (described in (40)) in Table 2 and, as predicted by constraints confining the atom movements, all lattice-constrained potentials were slightly stiffer. In addition, the GDP-tubulin potentials were stiffer than the GTP-tubulin potentials. Since the potential well-depths were not statistically different between the constrained and unconstrained cases (p>0.9), we calculated a mean longitudinal PMF for GDP- and GTP-tubulin, averaging all 15 replicates of constrained and unconstrained simulations together (Fig S6). To test whether different nucleotide states had different potential minima after the averaging, we performed a statistical analysis of the minima locations. In the same simulation configuration, i.e. constrained or unconstrained, none of the potential minima locations for either nucleotide state was significantly different (p>0.8), indicating that the GDP lattice compaction had relaxed into the GTP extended lattice. Overall, we conclude that the stronger longitudinal bond strength of GTP-tubulin compared to GDP-tubulin can explain why GTP-capped protofilaments would favor polymerization while GDP protofilaments would dissociate faster from a protofilament end due to weaker longitudinal bond strength.

### Brownian Dynamics Simulations Imply a Free Energy Difference, ΔΔG^0^_long_, of ≈ 4 k_B_T Between Nucleotide States

To predict the influence of the PMF on the subunit addition-loss kinetics and thermodynamics as quantified by the standard Gibbs free energy difference between the GTP and GDP nucleotide states, we performed BD simulations to model dimers’ diffusion and assembly kinetics on time scales up to 0.5 seconds, instead of only ~1μs via MD. Following our multi-scale approach, we calculated the full entropic penalty of longitudinal binding, which allows us to account for the fact that the stronger longitudinal bond (~25-32 k_B_T well depth) will have a higher entropic penalty compared to the weaker lateral bond (~11-12 k_B_T well depth; Hemmat *et al.*, 2019). MD entropy-corrected potentials (Fig 5A) were used as input to our BD simulations of microtubule lattice assembly (Fig 5B, 5C). Binding and unbinding of an incoming tubulin dimer to the tip of a protofilament with zero or one lateral neighbor were then simulated. Our zero-neighbor simulation results, as depicted in Table 3 show that a longitudinal energetic difference 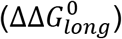 of ~3.5 ± 0.5 k_B_T exists between GDP- and GTP-states using the different input potentials obtained from the MD simulations for unconstrained dimers. To test the hypothesis that the slightly stiffer potentials from constrained MD simulations would change the BD model outputs, we ran another set of BD simulations using lattice constrained MD inputs. While the energetic difference mean value is higher for the constrained case, 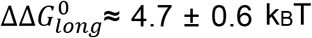, it is not statistically different from the unconstrained value (p > 0.6). We conclude that GTP tubulins have stronger longitudinal bonds as measured by 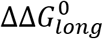 than GDP tubulins by 3.5 to 4.7 (≈4 on average) k_B_T.

**Table 3.** BD simulation results for dimer incorporation into a protofilament with zero lateral neighbors

| **Longitudinal bond** |  |  |  |  |
| --- | --- | --- | --- | --- |
| **Model estimated parameters** | **Lateral neighbors=0** |  |  |  |
|  | **Unconstrained** |  | **Constrained** |  |
|  | **GDP** | **GTP** | **GDP** | **GTP** |
| $k_{on}$ ($\mu\text{M}^{-1}\text{s}^{-1}\text{PF}^{-1}$) | $12.0 \pm 0.5$ | $35.5 \pm 2.0$ | $7.0 \pm 0.7$ | $41.4 \pm 1.9$ |
| $k_{off}$ ($\text{s}^{-1}$) | $(45.3 \pm 21.2) \times 10^3$ | $(4.0 \pm 1.2) \times 10^3$ | $(54.8 \pm 20.6) \times 10^3$ | $(3.1 \pm 1.6) \times 10^3$ |
| $\Delta G_B^0$ ($k_B T$) | -21.2 | -27.8 | -21.6 | -28.8 |
| $\Delta G_{long}^0$ ($k_B T$) | -5.6 | -9.1 | -4.9 | -9.5 |
| $\Delta G_s^0$ ($k_B T$) | +15.6 | +18.8 | +16.8 | +19.2 |
| $\Delta \Delta G_{long}^0$ ($k_B T$) | $3.5 \pm 0.5$ | | $4.7 \pm 0.6$ | |

PF<sup>-1</sup> = per protofilament

**Figure 5.**
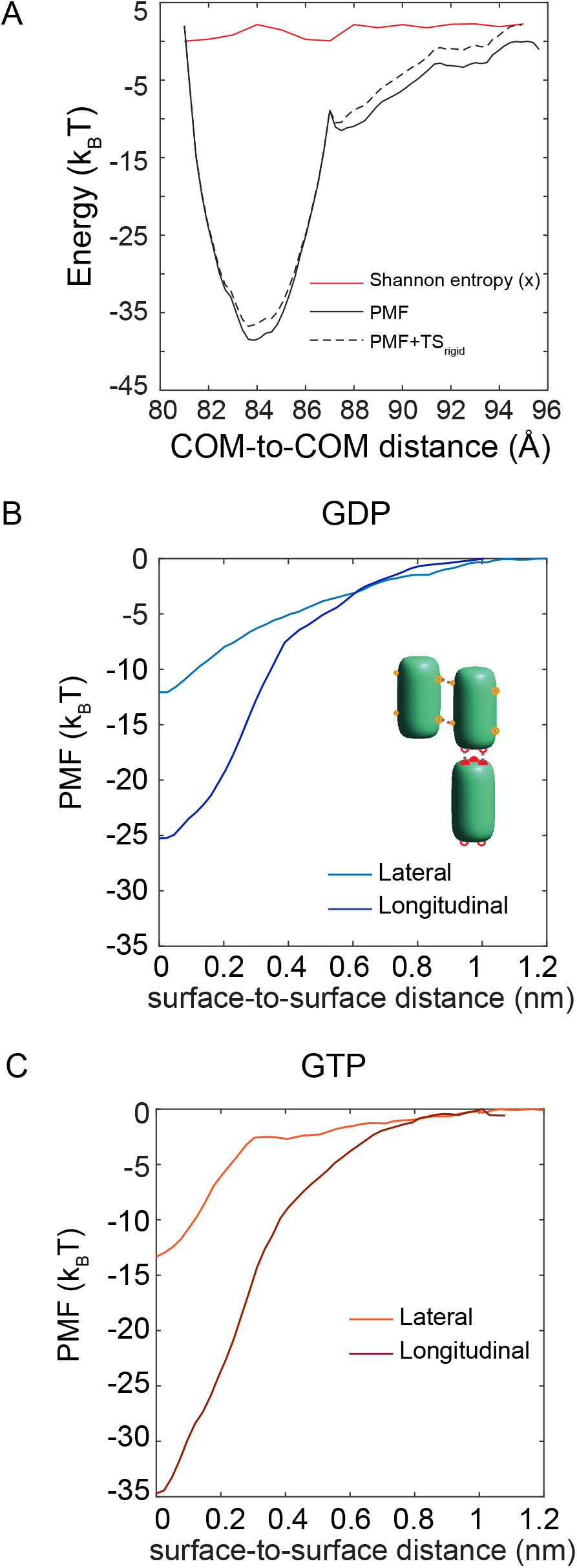
Longitudinal potentials derived from MD simulations used as inputs for BD simulations. (A) A replicate of PMF, its rigid-body Shannon entropy, and the entropy-corrected PMF (PMF+TS_rigid_) are shown for GTP-tubulin as a function of dimers’ distance. (B, C) BD lateral and longitudinal potential inputs are shown as a function of surface-to-surface distance for GDP- and GTP-tubulins respectively. Dark blue and orange show longitudinal potentials for GDP- and GTP-state, respectively, and light blue and orange indicate lateral potentials for GDP- and GTP-state, respectively.

To account for the stabilizing effect of a lateral neighbor, we ran BD simulations of dimer incorporation into a “cozy corner,” i.e. a protofilament tip with a longitudinal and a single lateral bond, using our lateral (40) and longitudinal potentials as inputs. The results, summarized in Table 4, yield the estimated lateral bond value for GDP- and GTP-states in the microtubule lattice, from comparing the total energy (∆*G*^0^) value to the zero-neighbor case. The lateral bond strength, as expected from the lateral potential inputs from our previous study (40), is found to be nucleotide independent, based on the values for unconstrained dimers 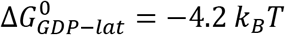 and 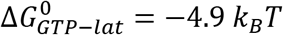, and 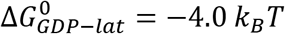 and 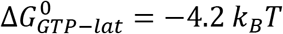 for constrained dimers, which are not statistically different (p > 0.7). In addition, the association rate constants, k_on_, and dissociation rate constants, k_off_, of GDP-tubulin are generally slower for the constrained input potentials as compared to the unconstrained input potentials. This is because as the potential becomes stiffer, it becomes harder for the dimer to diffuse in and out of the potential well (45). However, for GTP-tubulin, a significant change in the kinetics is not expected as the stiffness of the longitudinal potential is not significantly different in constrained vs. unconstrained simulations. Based on our previous BD simulations of bimolecular association in crowded environments, we do not expect the kinetics and thermodynamics to be significantly altered in *in vivo* environments due solely to macromolecular crowding (49).

**Table 4.** BD simulation results for dimer incorporation into a protofilament with one lateral neighbor

| **Longitudinal + Lateral bond** |  |  |  |  |
| --- | --- | --- | --- | --- |
| **Model estimated parameters** | **Lateral neighbors=1** |  |  |  |
|  | **Unconstrained** |  | **Constrained** |  |
|  | **GDP** | **GTP** | **GDP** | **GTP** |
| $k_{on}$ ($\mu\text{M}^{-1}\text{s}^{-1}\text{PF}^{-1}$) | $7.8 \pm 0.4$ | $23.2 \pm 1.1$ | $5.1 \pm 0.6$ | $27.8 \pm 1.6$ |
| $k_{off}$ ($\text{s}^{-1}$) | 447.6 | 20.2 | 702.8 | 30.0 |
| $\Delta G_B^0$ ($k_B T$) | -30.3 | -40.5 | -29.6 | -43.2 |
| $\Delta G_{tot}^0$ ($k_B T$) | -9.8 | -13.9 | -8.8 | -13.6 |
| $\Delta G_s^0$ ($k_B T$) | +20.5 | +26.5 | +20.7 | +29.5 |
| $\Delta G_{lat}^0$ ($k_B T$) | -4.2 | -4.9 | -4.0 | -4.2 |

PF<sup>-1</sup> = per protofilament

Altogether, these BD results give us insight into the possible mechanisms by which GDP-tubulin differs from GTP-tubulin in its stabilizing behavior in microtubule assembly that can be used in larger scale thermo-kinetic modeling of microtubule dynamic behavior. Specifically, these results are consistent with a model where a potential well-depth difference (∆*U*_*long*_) of ~6.6 k_B_T in the input longitudinal potential creates a 3.5-to 5-fold decrease in the on-rate constant, and an 11-to 22-fold increase in the off-rate constant of GDP-tubulin compared to GTP-tubulin, depending on their lateral neighbor case.

### Thermokinetic modeling identifies a preferred bending angle difference to reproduce experimental microtubule tip structures

Collectively, the MD and BD simulations show that GDP-tubulin pays an energetic penalty through its weaker longitudinal bond while the lateral bond strength remains nucleotide-independent. These results led us to hypothesize that the energetic penalty due to GTP hydrolysis in the microtubule lattice must either exist on the longitudinal bond only or on both longitudinal bond and bending flexibility and/or angle preference. Unfortunately, we do not yet have detailed structural information regarding the bending flexibility or angle preference of the dimers, and previous studies have not reported a potential for various bending modes of tubulin. To eliminate one of our possible hypotheses for dynamic instability, we ran our previously described thermokinetic model (34,38,40) (Fig 6A) with ΔΔG^0^ implemented on the longitudinal bond alone or in combination with lateral bond energetic differences due to possible nucleotide-dependent bending flexibility/mechanics, which allowed us to predict microtubule net assembly rates, and tip structures during net polymerization and depolymerization. We first ran a single state thermokinetic model of microtubule assembly using our BD estimated on rate constants and *in vitro* parameter set (Table S4). The heatmap of the resultant microtubule net assembly rates (Fig 6B) indicates that our MD-BD estimated free energies are in reasonable agreement with published in vitro assembly rates (50).

**Figure 6.**
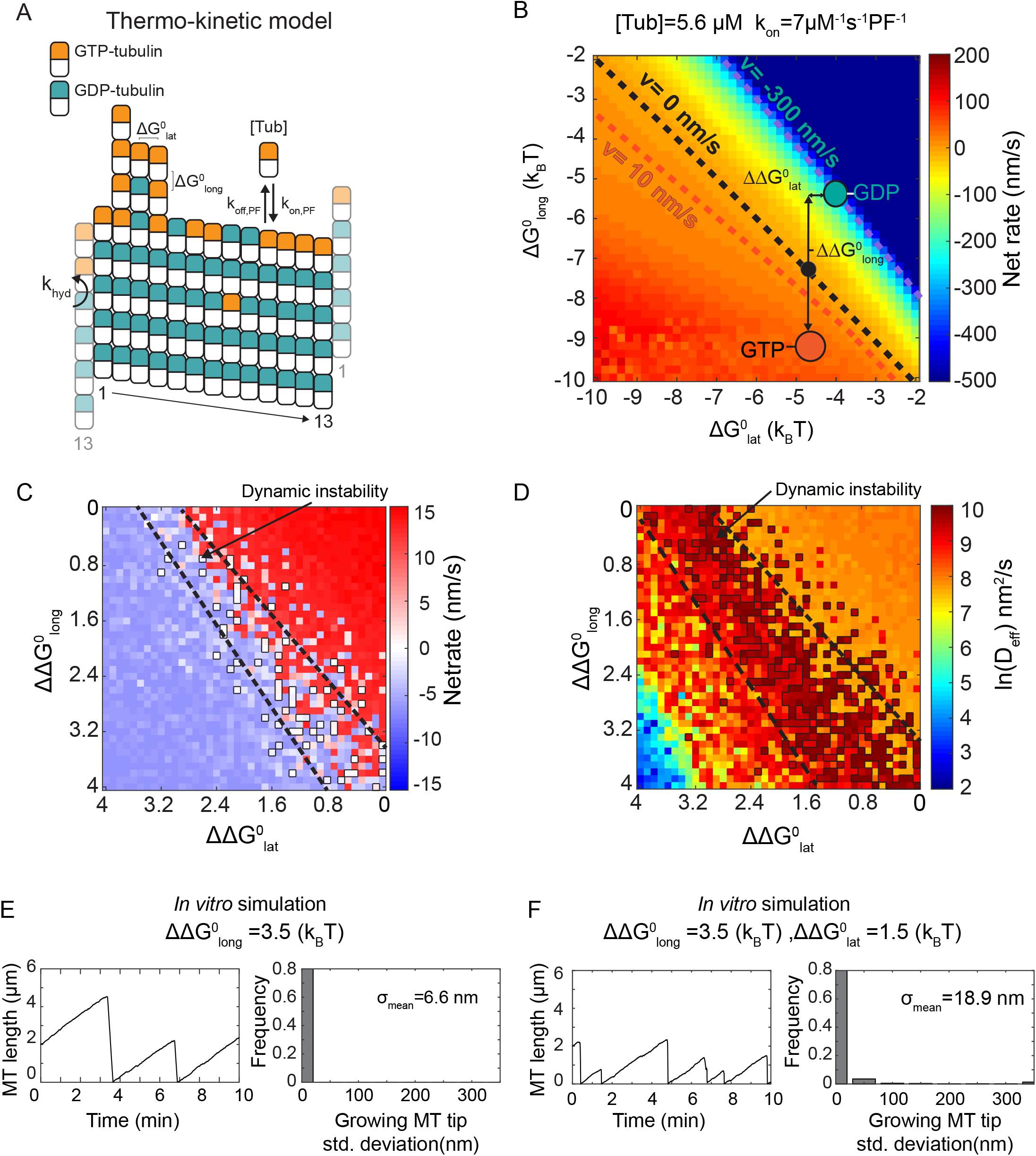
Thermokinetic model for microtubule self-assembly to capture *in vitro* dynamics. (A) Base thermokinetic model with its parameters, as described in VanBuren *et al.* 2002, Gardner *et al.*, 2011, Castle *et al.* 2013. (B) Microtubule net assembly rates shown as a function of different lateral 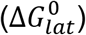 and longitudinal 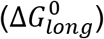 bond free energies using the *in vitro* parameter set (Table S4). The net-rates are obtained using a pseudo-mechanical model without hydrolysis. Blue and orange circles show our predicted areas where GDP- and GTP-tubulin are located, respectively. Black circle shows the reference point for dynamic instability. Lateral and longitudinal energy penalties are shown as 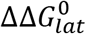 and 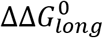 respectively. (C) Microtubule net-rate values as a function of varying lateral and longitudinal energetic penalties. Black outlined regions, between the dashed line, show zero-net rates where dynamic instability is observed. (D) Microtubule apparent diffusion coefficient derived from microtubule length variance shown as a function of varying lateral and longitudinal energetic penalties. Black outlined regions, between the dashed lines, show diffusion values consistent with *in vitro* experiments. (E) Microtubule length vs. time and associated tip standard deviation distribution for an energetic penalty on longitudinal bond only, 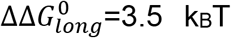, or (F) on both the longitudinal and lateral bonds, 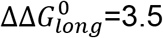, 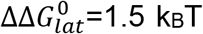. Mean standard deviation of the tip, σ_mean_, is indicated on the plots.

Based on our previous analyses, microtubules undergo dynamic instability with zero net rate overall (averaged over multiple rounds of dynamic instability) with an apparent microtubule tip net assembly diffusion coefficient of ~0.8-16 × 10^4^ nm^2^/s, extracted from experimental microtubule length variance measurements (38,50). As shown in Fig 6C and 6D, the calculated microtubule net assembly rates and apparent diffusion coefficients as a function of ΔΔG_lat_0 and ΔΔG_long_0 are consistent with those reproducing dynamic instability (highlighted inside the dashed line borders). Dynamic instability is found over a range of lateral energetic penalties of less than 1.5 k_B_T combined with the MD-BD estimated ΔΔG_long_0 of 3.5 ± 0.5 k_B_T. Interestingly, if the energetic penalty of hydrolysis that destabilizes the lattice is only paid through the longitudinal bond, microtubules would grow and shorten with protofilaments having very low protofilament length standard deviation, σ_tip_ (Fig 6E), i.e. “blunt” tips (while a larger σ_tip_ reflects “tapered” tips) (50,51). However, if the penalty is applied to both longitudinal and bending angle preference and/or flexibility (i.e. the lateral bond in our model), tapered tips with variable protofilament lengths and assembly dynamics appear in the simulated microtubules, manifested as a larger σ_tip_ (Fig 6F). This finding is consistent with previous *in vitro* (50) and *in vivo* (51) experiments that reported highly variable tapered tips for microtubules by high resolution imaging and microtubule tip tracking. It is also in line with previous work that suggested there is a flexibility difference between the two nucleotide-states, with GTP being softer at its intra- and inter-dimer interface (29,35,41), or a bending preference (24,33) with GTP-protofilaments growing less curved or nearly straight compared to the extensive outward peeling observed for GDP-protofilaments.

### Mechanochemical modeling indicates an outward bending preference is required to recreate both dynamic instability and microtubule tip structures

Since both bending flexibility and angle preference are directly related to mechanics of a microtubule lattice, using a more detailed model where mechanics are directly modeled can help to pinpoint the possible mechanisms of dynamic assembly. The previously developed mechanochemical model of microtubule assembly (33) was used in our study to investigate whether a difference in bending angle preference (in both radial and tangential directions) and/or in bending flexibility combined with a ΔΔG^0^_long_ will recreate experimentally observed assembly rates as well as tip structures. We first tested the possibility of GDP-tubulin having higher radially outward bending preference (θ_x_^GDP^=22°, chosen according to (33), Fig 7A). The results show microtubule dynamic instability (Fig S7A), blunt growing microtubules (Fig S8A) and three major types of shortening tips: blunt, splayed and tapered (Fig 7A), consistent with experimentally observed EM tip structures (52,53). Tangential bending preference was then investigated as a possible mechanism by itself (Fig 7B) or combined with radially bending kink (Fig 7C). Our model showed that dynamic instability is disrupted, microtubules rarely undergo catastrophe, and mostly grow in the case of a tangential bending kink only, with θ_y_ ranging from 11°-22° (Fig S7B, S8B). However, combined with radial outward bending, microtubules show dynamic behavior (Fig S7C, S8C) and tip structures similar to Fig 7A, meaning that a combination of bending preference both radially and tangentially is plausible for microtubule assembly.

**Figure 7.**
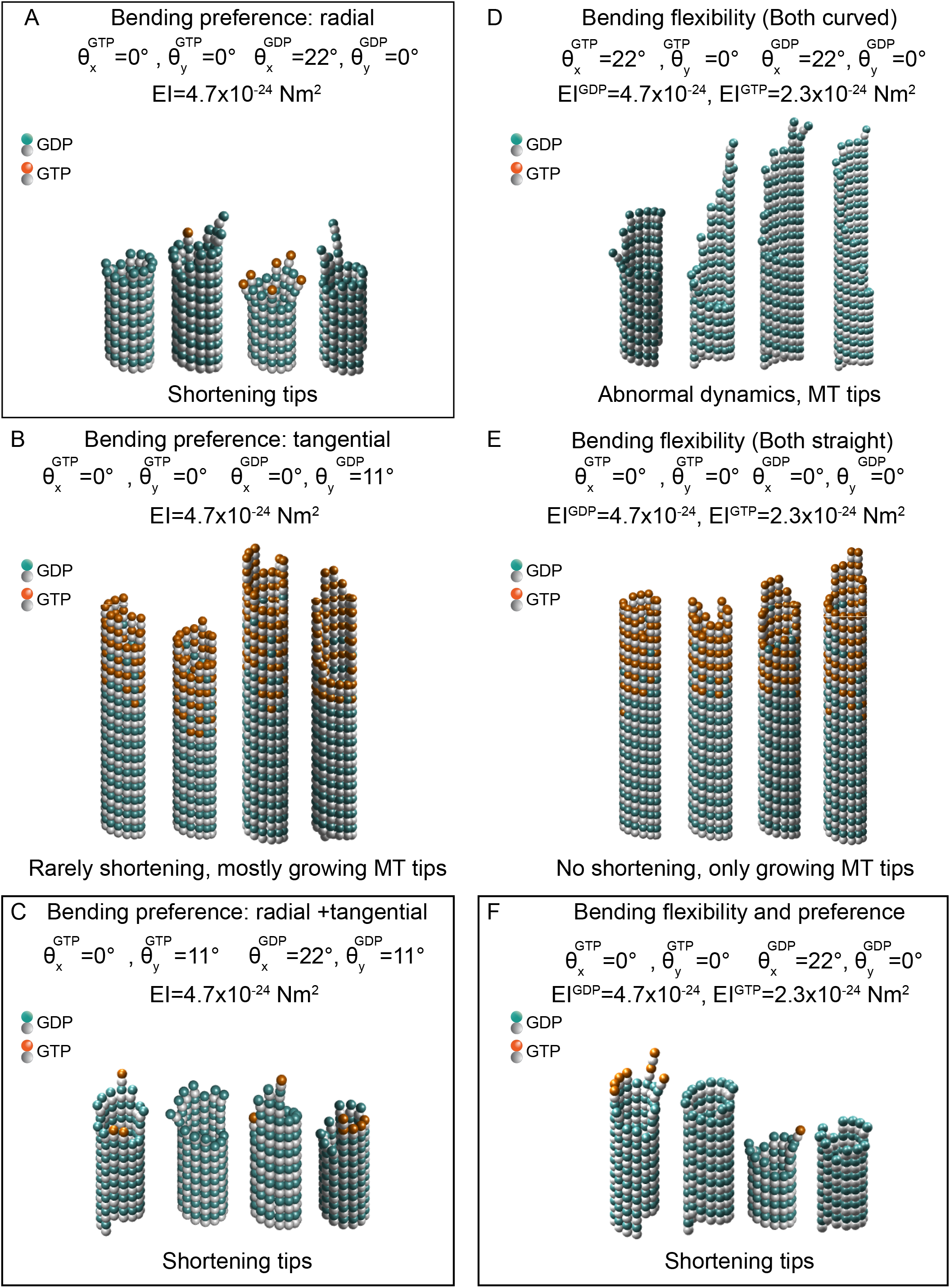
Simulated shortening microtubule tip structures using the mechanochemical model of microtubule assembly indicate possible mechanisms for dynamic instability. Shortening microtubules start from a blunt uncapped configuration and are run for 500 events. (A) Shortening microtubule tip structures for GDP-tubulin having a preferred radially outward bending angle (θ_x_^GDP^) of 22°, and GTP-tubulin with θ_x_^GTP^=0°, both having the same dimer flexibility. Shortening tip structures are either blunt, splayed, or tapered. (B) Microtubule tip structures for GDP-tubulin having a preferred tangential bending angle (θ_y_^GDP^) of 11°, and GTP-tubulin with θ_y_^GTP^=0°, both having the same dimer flexibility. Microtubules shorten very rarely and only blunt or tapered growing microtubule tip structures are observed. (C) Shortening microtubule tip structures for GDP-tubulin having θ_x_^GDP^=22°, and GTP-tubulin with θ_x_^GTP^=0°, both having the same dimer flexibility and tangential bending preference, θ_y_^GDP^= θ_y_^GTP^ =11°. Shortening tip structures are either blunt, splayed, or tapered. (D) Microtubule tip structures for GTP-tubulin having higher flexibility than GDP-tubulin, as EI^GTP^=2.3×10^−24^, EI^GDP^=4.7×10^−24^ Nm^2^, both having the same radial bending preference as curved, θ_x_^GDP^= θ_x_^GTP^ =22°. Normal microtubule dynamics were not observed. (E) Microtubule tip structures for GTP-tubulin having higher flexibility than GDP-tubulin, as EI^GTP^=2.3×10^−24^, EI^GDP^=4.7×10^−24^, both having the same radial bending preference as straight, θ_x_^GDP^= θ_x_^GTP^ =0°. Microtubule shrinkage is not observed. Blunt or tapered growing tip structures are shown. (F) Microtubule tip structures for GTP-tubulin having higher flexibility than GDP-tubulin, as EI^GTP^=2.3×10^−24^, EI^GDP^=4.7×10^−24^ Nm^2^, and GDP-tubulin having a preferred radially outward bending preference, θ_x_^GDP^=22°, θ_x_^GTP^ =0°. Shortening tip structures are either blunt, splayed or tapered. Bold borders show the mechanisms consistent with experimental results.

Next, as argued in previous studies (29,35), we assessed the possibility of a flexibility difference between different nucleotide states as the main cause of dynamic instability (Fig 7D, 7E). Interestingly, assuming that GTP-dimer is more flexible than GDP-dimer (~ 2 fold based on Table S1A) and both dimers prefer a radially kinked or straight conformation (θ_x_=22°, 0°) did not result in normal microtubule dynamics, i.e. abrupt microtubule length changes with extended periods of dynamic pause for both dimers kinked, and microtubules growing consistently without any shrinkage for both dimers preferring to be straight (Fig S7D, S7E, S8D, S8E). Lastly, a combination of flexibility difference along with a differential radial bending preference was applied to our model. Dynamic instability was observed (Fig S7F, S8F) and shortening microtubules showed either splayed, blunt or tapered tip structures (similar to Fig 7A, 7C).

These results highlight the importance of a radial outward bending angle preference for GDP-tubulin that exceeds that of GTP-tubulin in mediating microtubule dynamic assembly and tip structures. Although we cannot rule out that GDP-tubulin’s radial bending preference might be accompanied by other mechanisms such as flexibility differences or tangential bending preference as well, we conclude that radially outward kinking of GDP-tubulin is necessary for the model to agree with the experimental dynamics and tip structures.

## Discussion

Here we investigated the fundamental atomistic and molecular mechanics underlying a complex biological phenomenon, microtubule dynamic instability, by using a multiscale approach, integrating structural, mechanochemical, and kinetic perspectives that span from atoms to cellular scales. Our MD results, using previously published structures of tubulin, show that the longitudinal bond, in contrast to the lateral bond, is nucleotide dependent and is ~3 times stronger than lateral bond. However, we find that a longitudinal bond difference is insufficient by itself to produce the experimentally observed tapered growing tip structures, and a nucleotide-dependent radial preferred angle is essential to recreate curling protofilaments commonly found at the tips of shortening microtubules and blunt tip structures in growing microtubules. Thus, by using this multiscale approach without parameter adjustment, we conclude that dynamic instability occurs primarily by weakening of the longitudinal bond (~4 k_B_T) and secondarily by outward curling between dimers (~1.5 k_B_T) upon GTP hydrolysis in the microtubule lattice. More generally, these results show how dynamic simulations can be used to leverage atomistic structural data to identify the mechanistic origins of cell-level dynamics that are important to cell behavior.

Although the possibility of a nucleotide-regulated bending flexibility has received much attention recently (29,35,54,55), the suggested mechanism did not reproduce microtubule dynamics and predicted tip structures within our experimentally-constrained approach. An implication of this finding is that there is another important factor playing a role in maintaining dynamic instability, as observed in experiments. Nucleotide-dependent lateral and longitudinal bonds, with GDP-tubulin having a stronger longitudinal and a weaker lateral bond, suggested by a recent cryo-EM study (32) were also ruled out based on the results of multiscale model and our previous MD studies of the lateral bond PMF (37,40). It was only the addition of a radial bending preference to our nucleotide-dependent longitudinal bond in our model that captured both predicted microtubule dynamics and tip structures. Our results agree with the findings of Rice *et al.* (2008) where soluble dimers have similar intradimer conformations in solution, and further we show that the intradimer bending angles remain small and nucleotide-independent upon lattice constraints. In addition, in the presence of the lattice constraints, an outward radial interdimer bending preference (≈ 22°) along the protofilament is modulated by the nucleotide state. As an alternate mechanism, a GTP-dimer with a stronger longitudinal bond, a weaker radially outward bending angle and higher dimer flexibility cannot be ruled out within our methodology, consistent with experimental boundaries.

Despite the fact that our results are in good agreement with experimental microtubule dynamic assembly measurements and previously estimated microtubule computational models (33,34), it could be argued that our MD estimated potential energies can be affected both by the initial crystal structures, and by the selected reaction coordinate. The initial cryo-EM structures used in our study were obtained in the presence of kinesin, a tubulin dimer marker (37), which has been speculated to affect GDP-tubulin longitudinal compaction (56), or GMPCPP-lattice spacing (57). We note that although longitudinal compaction in the lattice might be regulated by kinesin to a degree, the instant compaction release and plateauing after 100ns in our MD simulations in the absence of kinesin justifies that any external effect is quickly relaxed (Fig 2, Fig S4). The fact that we obtain consistent PMFs in both constrained and unconstrained simulations further shows that our PMF calculation is reliable within the replicate-to-replicate variability (± 1.8 to 3.5 k_B_T SEM) and is not dependent on the initial conditions. Since we were unable to investigate all possible reaction coordinates, we chose the most probable distance-based path to investigate the bond energies between the dimers as determined by BD simulation (Fig S5).

The predictions made by our multiscale modeling approach show that although microtubule assembly dynamics have multiple parameters governing their behavior in the thermokinetic model, it is tightly regulated and only a limited range of parameters would agree well with the experimental results. Having a multiscale approach is significantly beneficial in narrowing down the predictions made from atomic/molecular level to cell-level behavior. This multiscale methodology can be a framework to predict the dynamics of other self-assembled polymers, especially those whose subunits are relatively rigid and ordered, such as F-actin, deoxygenated sickle hemoglobin fibers, amyloid fibrils, and virus capsids to make physiologically relevant predictions from atomic changes, such as the effects of hydrolysis, mutations, post-translational modifications, microtubule-associated proteins, drugs, and therapeutic agents.

## Methods

### Molecular Dynamics Simulations

MD Simulations of all systems were run using NAMD 2.10 software package (58) using the CHARMM 36 force field (59) for parametrization. VMD 1.9 (60) was used for visualization and trajectory analysis. Longitudinally-paired tubulins, both dimers with GDP-or GMPCPP-nucleotides, were extracted from the published cryo-EM structures of microtubules by Zhang et al. (37) (PDB ID 3JAS, 3JAT). GTP-tubulin structure was built based on GMPCPP-tubulin structure by swapping GMPCPP out for a GTP, which was necessitated by lack of a true GTP-tubulin structure in a microtubule lattice. The protein complex along with the nucleotides were all parametrized using the CHARMM-GUI interface (61). Each simulation system was initially energy minimized for 12000 steps using the conjugate gradient algorithm, and then they were solvated in TIP3P water (62), using a 10 Å margin from each side. The simulation systems were neutralized with MgCl_2_ ions at 2mM concentration based on physiologically-relevant salt concentrations.

The solvated systems were heated to 310 K for 1ns using a Langevin thermostat (63), and then run in an NPT ensemble (T=310K and P=1 atm). The simulations were followed by a total production run of 350ns for each system (after 50ns equilibration). All simulations were run with 2 fs time step and a cutoff radius of 12Å for van der Waals interactions, using Particle Mesh Ewald (PME) for long range non-bonded interactions (64). The equilibrium run trajectories were stored every 3000-time steps (6 ps). RMSD, RMSF, hydrogen bonds and salt bridges were calculated using manually written tcl scripts and plugins available in VMD. The buried solvent-accessible surface area was calculated as previously described (40).

To simulate the effect of lateral neighbors in the microtubule lattice without increasing the number of atoms, we identified lateral bond residues from (40) and longitudinal bond residues from our unconstrained simulations and applied a harmonic constraint to those atoms. The stiffness of the constraints of lateral and longitudinal neighbors were chosen according to the stiffness of the lateral PMF (40), *κ*_*lat*_ = 1 *kcal*/*mol* Å^2^ and unconstrained longitudinal PMF, *κ*_*long*_ = 1 *kcal*/*mol* Å^2^.

For free energy calculations, we employed the umbrella sampling method (65) combined with weighted histogram analysis method (WHAM) (47,48) to be able to sample the ensemble sufficiently and have independent simulations that each can be run for longer sampling time in parallel, considering the large number of atoms.

A PMF, a free energy landscape as a function of a specified reaction coordinate, was obtained for each nucleotide case. The reaction coordinate was defined as the longitudinal COM to COM distance of the dimers, without any rotations, since that is the most probable path of longitudinal unbinding of the dimers according to our BD simulations (Fig S5). The bias potential stiffness was tuned to be 10 kcal mol^−1^Å^−2^ to give sufficient overlap of the histograms of the windows. Fifteen windows were created, each being 1Å separated from their nearest window. The reaction coordinate in free energy simulations was recorded every 200-time steps (0.4 ps).

To ensure the time convergence of the PMFs to the ensemble-average PMFs, we employed similar methodology in (40) to run multiple replicates with different initial conditions and to gradually increase window sampling time until the PMF change did not exceed a threshold of 1.5 k_B_T, determined by the Monte Carlo bootstrap error of the PMFs. A window sampling time of 30ns was sufficient to produce a convergence in each PMF. The last 200ns of the equilibrium run was used to choose equilibrated initial structures for creating replicates of umbrella windows for each system.

NVIDIA Tesla K40 GPUs were used to accelerate the simulations on the Mesabi cluster at the Minnesota Supercomputing Institute (MSI), University of Minnesota, and NVIDIA Kepler K80 GPUs were used on Comet and Bridges, Extreme Science and Engineering Discovery Environment (XSEDE) (66) dedicated clusters at the San Diego Supercomputing Center (SDSC) and Pittsburgh Supercomputing center (PSC).

### Brownian Dynamics Simulations

Tubulin dimers’ association and dissociation from a protofilament in microtubule lattice was simulated using the BD model of Castle *et al.* (45). The entropy corrected PMFs, (Fig 5A), with entropy calculated based on the rigid body Shannon entropy method as described in (40), were used as the input to the BD simulations. Simulations were run for two nucleotide cases and each was run for a total of 500,000 iterations of binding simulations and 20 to 20,000 iterations of unbinding simulations (total time of 0.1-1s). For binding simulations, half-force radius was calculated for the PMF energy profiles and used as the binding radius in BD simulations (40). For unbinding simulations, we used a separation distance criterion of R_U_=11 nm, according to (45), where the probability of rebinding is very low (p <0.01). BD of dimer’s dissociation from a protofilament with two lateral neighbors was not simulated due to extremely high stability and long unbinding time (no unbinding event up to 0.1s).

### Thermokinetic and Mechanochemical Modeling

Microtubule assembly dynamics were simulated using a pseudo-mechanical thermo-kinetic modeling, as previously developed (34) and modified (38). For all simulations, an *in vitro* parameter set was used (Table S4) with variable energy penalty for hydrolysis. Seed length and starting GTP layers were set to 2 µm. On-rate penalties of 2 and 10 were added for one and two lateral neighbor cases, respectively (45). Microtubule tip structures were obtained using the mechanochemical model as previously described (33). All the shortening microtubules were simulated starting from uncapped protofilaments with blunt tips, ran for 500 events. Model parameters were similar to (33), with modification of preferred bending angles and flexibilities. Higher flexibility in GTP-dimer was chosen according to bending angle data variance (Table S1A) as double the flexibility of GDP-dimer. As for bending preference, the preferred radial bending angle was selected as 22° (33), and the preferred tangential bending angle was set to 11°, according to Table S1A, where the tangential mean value is almost half of the radial mean value.

## Author Contributions

Conceptualization, M.H. and D.J.O.; Methodology, M.H., and D.J.O.; Software, M.H.; Analysis, M.H., and D.J.O.; Writing – Original Draft, M.H. and D.J.O.; Writing – Review & Editing, all authors; Visualization, M.H.; Supervision, D.J.O.; and Funding Acquisition, D.J.O.

## Acknowledgements

The authors thank Dr. Jonathan Sachs for advice and helpful discussions. This study was supported by National Institutes of Health under award number R01-AG053951 and the Institute for Engineering in Medicine (IEM) award at the University of Minnesota to DJO. The authors acknowledge the Extreme Science and Engineering Discovery Environment (XSEDE), Comet system at the San Diego Supercomputing Center (SDSC) and Bridges system at the Pittsburgh Supercomputing Center (PSC) through allocation MCB160060, and the Minnesota Supercomputing Institute (MSI) at the University of Minnesota for providing resources that contributed to the research results reported within this paper.

