## Supplementary Material and figures for "Atomistic basis of microtubule dynamic instability assessed via multiscale modeling"

### Supporting Methods

#### Tubulin bending angle and flexibility analysis

Tubulin dimer bending angles were calculated based on the methodology described in (1), similar to (2, 3). Two bending motions (radial and tangential to microtubule wall) were identified throughout constrained and unconstrained simulations for both nucleotide states (Fig S1A, S1B). To further investigate the effect of the lattice on the flexibility of the dimers, we calculated the root-mean-square fluctuation (RMSF) of the backbone residues averaged over the last 300ns of the trajectories, for GDP- and GTP-tubulin (Fig S1C, S1D). As indicated in the bending motion, when we get farther from the lattice constraints, motion fluctuations of the residues are higher on average (red color in the RMSF map). When the RMSFs of the two nucleotide states are compared for each residue (Fig S1E, S1F), the lattice constraints confine GTP-tubulin dimer motions to a higher degree as compared to GDP-tubulin, while the free GTP-dimer fluctuates more than GDP-tubulin.

For intradimer bending angle calculation, α-tubulin subunit of the dimer was first aligned and then the angle between the vector connecting the COMs of α- and β-subunits and the reference vector was calculated ($\theta_{dot}$). For interdimer bending angle, we employed a similar method to the one applied to the intradimer interface, with the vectors connecting COM of the first dimer to the second dimer. We further decomposed the angles to 3 perpendicular components to identify the dominant direction of bending and their autocorrelation times, as seen in Fig S2 and S3 for the unconstrained and constrained dimers.

To compare the average values and distribution of the angles for the two nucleotide states, we calculated the mean of 100 bootstrapped data points, selected far apart (twice the autocorrelation characteristic time) to avoid correlation, from the last 300ns of the trajectories, summarized in Table S1A, S1B, S2A and S2B. To characterize the angle fluctuations magnitude, flexural stiffness was calculated based on the equipartition theorem (4), where thermal motion variance is inversely related the flexural stiffness:

$$\kappa = \frac{k_B T}{\sigma^2} \quad (1)$$

Where $k_B$ is Boltzmann's constant, T is absolute temperature, and $\sigma^2$ is the variance. The autocorrelation function is calculated by *autocorr* function in MATLAB, and the characteristic time ($\tau_{ACF}$) is obtained via fitting an exponential decay function.

We believe that these bending motions are strongly auto-correlated (Table S1B, Table S2B) and highly variable among different simulation replicates (as observed in Fedorov *et al.*'s Fig 3C, 3D, 4C, and 4D in (5)). The bending angles for both GDP- and GTP-tubulins showed that, compared to the unconstrained simulations, the tangential and radial bending motions of the constrained dimers are limited, and they maintain a straighter conformation, as expected (Fig S1B, S3A, S3B). We found that the intradimer angles' fluctuations for the bottom dimer are significantly decreased due the lattice constraints, while for the top dimer the lattice constraint effects are less significant (Fig

S3B). Interestingly, we found that GTP-dimer bending motions are more confined in the lattice than GDP-dimer, whereas at the protofilament tip where there are no neighbor constraints, GTP-tubulin exhibits a higher range of motion (Table S2A). This result suggests that the presence of the microtubule lattice modulates bending stiffnesses.

**Optimization of umbrella sampling method**

The collective variable sampled in our umbrella sampling method was chosen as the COM-to-COM distance of the two dimers. To ensure that our reaction path is the most probable path taken by the dimers to bind to each other, we ran BD simulations of separate dimers, initially located randomly in a sphere of 1.5 nm radius, binding to each other and inspected the binding efficiency as a function of the initial relative rotation angles (Fig S5). The results show that only a limited range of angles (<4 degrees) has a high probability of binding, meaning that separating the dimers longitudinally without a major rotation is not an unreasonable choice of the reaction coordinate.

### Supporting Tables

#### Table S1A. Analysis of interdimer and intradimer bending angles of unconstrained tubulin dimers in GDP- and GTP-state

| Simulation | Interdimer angles | | | | | |
|---|---|---|---|---|---|---|
| | $\theta_x$ (Radial) | | $\theta_y$ (Tangential) | | $\theta_z$ (Twist) | |
| Nucleotide | Mean ± SEM (deg) | Flexural Rigidity ± SEM ($k_BT/rad^2$) | Mean ± SEM (deg) | Flexural Rigidity ± SEM ($k_BT/rad^2$) | Mean ± SEM (deg) | Flexural Rigidity ± SEM ($k_BT/rad^2$) |
| GDP | 8.9 ± 0.2 | 660.2 ± 83.9 | 6.8 ± 0.3 | 453.7 ± 73.0 | 0.6 ± 0.0 | (46.1 ± 7.5) x $10^3$ |
| GTP | 15.9 ± 0.3 | 311.1 ± 46.2 | 6.5 ± 0.3 | 495.8 ± 75.7 | 0.7 ± 0.0 | (19.3 ± 2.9) x $10^3$ |

| | Intradimer angles (Top dimer) | | | | | |
|---|---|---|---|---|---|---|
| Simulation | $\theta_x$ (Radial) | | $\theta_y$ (Tangential) | | $\theta_z$ (Twist) | |
| Nucleotide | Mean ± SEM (deg) | Flexural Rigidity ± SEM ($k_BT/rad^2$) | Mean ± SEM (deg) | Flexural Rigidity ± SEM ($k_BT/rad^2$) | Mean ± SEM (deg) | Flexural Rigidity ± SEM ($k_BT/rad^2$) |
| GDP | -0.1 ± 0.1 | 2317.8 ± 446.5 | -0.2 ± 0.1 | 3563.2 ± 484.4 | 0.0 ± 0.0 | (5.6 ± 1.2) x $10^6$ |
| GTP | 0.2 ± 0.1 | 1987.3 ± 341.0 | 3.8 ± 0.1 | 2576.2 ± 459.9 | 0.0 ± 0.0 | (3.6 ± 0.6) x $10^6$ |

#### Table S1B. Autocorrelation times of inter- and intradimer angle time series of unconstrained tubulin dimers in GDP- and GTP-state

| | Interdimer angles $\tau_{ACF}(ns)$ | | |
|---|---|---|---|
| Nucleotide | $\theta_x$ | $\theta_y$ | $\theta_z$ |
| GDP | 5.9 | 34.1 | 8.9 |
| GTP | 9.2 | 6.7 | 6.8 |
| | Intradimer angles (Top dimer) $\tau_{ACF}(ns)$ | | |
| Nucleotide | $\theta_x$ | $\theta_y$ | $\theta_z$ |
| GDP | 32.4 | 5.1 | 18.9 |
| GTP | 15.0 | 14.5 | 11.0 |

**Table S2A. Analysis of interdimer and intradimer bending angles of constrained tubulin dimers in GDP- and GTP-state**

| Simulation | Interdimer angles | | | | | |
|---|---|---|---|---|---|---|
| | $\theta_x$ (Radial) | | $\theta_y$ (Tangential) | | $\theta_z$ (Twist) | |
| Nucleotide | Mean ± SEM (deg) | Flexural Rigidity ± SEM ($k_BT/rad^2$) | Mean ± SEM (deg) | Flexural Rigidity ± SEM ($k_BT/rad^2$) | Mean ± SEM (deg) | Flexural Rigidity ± SEM ($k_BT/rad^2$) |
| GDP | 5.8 ± 0.2 | 1015.6 ± 127.3 | 2.8 ± 0.3 | 380.9 ± 75.9 | 0.6 ± 0.0 | (0.1 ± 0.0) x $10^6$ |
| GTP | 2.8 ± 0.2 | 1016.7 ± 158.2 | -0.3 ± 0.1 | 1451.5 ± 189.9 | 0.0 ± 0.0 | (3.6 ± 0.4) x $10^6$ |

| | Intradimer angles (Top dimer) | | | | | |
|---|---|---|---|---|---|---|
| Simulation | $\theta_x$ (Radial) | | $\theta_y$ (Tangential) | | $\theta_z$ (Twist) | |
| Nucleotide | Mean ± SEM (deg) | Flexural Rigidity ± SEM ($k_BT/rad^2$) | Mean ± SEM (deg) | Flexural Rigidity ± SEM ($k_BT/rad^2$) | Mean ± SEM (deg) | Flexural Rigidity ± SEM ($k_BT/rad^2$) |
| GDP | 2.0 ± 0.1 | 2751.4 ± 391.6 | 1.9 ± 0.1 | 1789.0 ± 316.8 | 0.0 ± 0.0 | (11.2 ± 2.2) x $10^6$ |
| GTP | 0.3 ± 0.1 | 2462.3 ± 512.3 | 4.4 ± 0.1 | 3883.7 ± 626.0 | 0.0 ± 0.0 | (2.4 ± 0.5) x $10^6$ |

**Table S2B. Autocorrelation times of inter- and intradimer angle time series of constrained tubulin dimers in GDP- and GTP-state**

| | Interdimer angles $\tau_{ACF}(ns)$ | | |
|---|---|---|---|
| Nucleotide | $\theta_x$ | $\theta_y$ | $\theta_z$ |
| GDP | 3.8 | 26.5 | 22.9 |
| GTP | 14.8 | 11.6 | 1.0 |
| | Intradimer angles (Top dimer) $\tau_{ACF}(ns)$ | | |
| Nucleotide | $\theta_x$ | $\theta_y$ | $\theta_z$ |
| GDP | 6.0 | 15.4 | 2.5 |
| GTP | 27.7 | 9.6 | 28.5 |

**Table S3.** Longitudinal potential well-depths for different replicates and nucleotides in the unconstrained and constrained simulations

| Replicate | Unconstrained 30ns sampling time* | | Constrained 30ns sampling time | |
|---|---|---|---|---|
| | $U_{GDP}$ ($k_B$T) | $U_{GTP}$ ($k_B$T) | $U_{GDP}$ ($k_B$T) | $U_{GTP}$ ($k_B$T) |
| 1 | 25.2 ± 1.4** | 33.5 ± 1.2 | 28.6 ± 1.3 | 37.3 ± 1.4 |
| 2 | 23.2 ± 1.4 | 22.0 ± 1.4 | 24.3 ± 1.4 | 27.4 ± 1.5 |
| 3 | 29.4 ± 1.4 | 38.1 ± 1.0 | 37.3 ± 1.3 | 36.2 ± 1.5 |
| 4 | 24.5 ± 1.0 | 36.5 ± 1.4 | 17.6 ± 1.3 | 29.2 ± 1.1 |
| 5 | 18.6 ± 0.9 | 31.5 ± 1.5 | 19.5 ± 1.5 | 34.2 ± 1.3 |
| 6 | 32.9 ± 1.4 | 32.3 ± 1.2 | N/A | N/A |
| 7 | 18.8 ± 1.5 | 26.4 ± 1.5 | N/A | N/A |
| 8 | 21.2 ± 1.5 | 39.6 ± 1.5 | N/A | N/A |
| 9 | 18.5 ± 1.5 | 27.8 ± 1.5 | N/A | N/A |
| 10 | 31.9 ± 1.5 | 31.1 ± 1.5 | N/A | N/A |

* Sampling time is chosen to be when the bootstrap Monte Carlo error drops below 1.5 $k_B$T

** All values are calculated based on the average of 10 points near the absolute minimum and maximum of the potential due to thermal fluctuations

**Table S4.** *In vitro* simulation base parameter set

| Parameter | Definition | Value | Reference |
|---|---|---|---|
| $\Delta G_{lat}^0$ | Lateral bond free energy | -5 $k_B$T | Hemmat *et al.* (2019); Gardner *et al.* (2011) |
| $\Delta G_{long}^0$ | Longitudinal bond free energy | -7.2 $k_B$T | Current work; Gardner *et al.* (2011) |
| $\Delta\Delta G^0$ | Energetic penalty of GDP-tubulin | +3.6 $k_B$T | Current work |
| [Tub] | Free tubulin concentration | 5.6 μM | Schek *et al.* (2007); Gardner *et al.* (2011) |
| $k_{on,PF}$ | On-rate constant | 6 $\mu M^{-1}$ $s^{-1}PF^{-1}$ | Gardner *et al.* (2011) |
| $k_{hyd}$ | Hydrolysis rate constant | 0.2 $s^{-1}$ | Model constrained; Coombes *et al.* (2013) |
| $\sigma_1$, $\sigma_2$ | One- and two-neighbor on-rate penalty | 2, 10 | Castle *et al.* (2013) |

PF = protofilament

### Supporting Figures

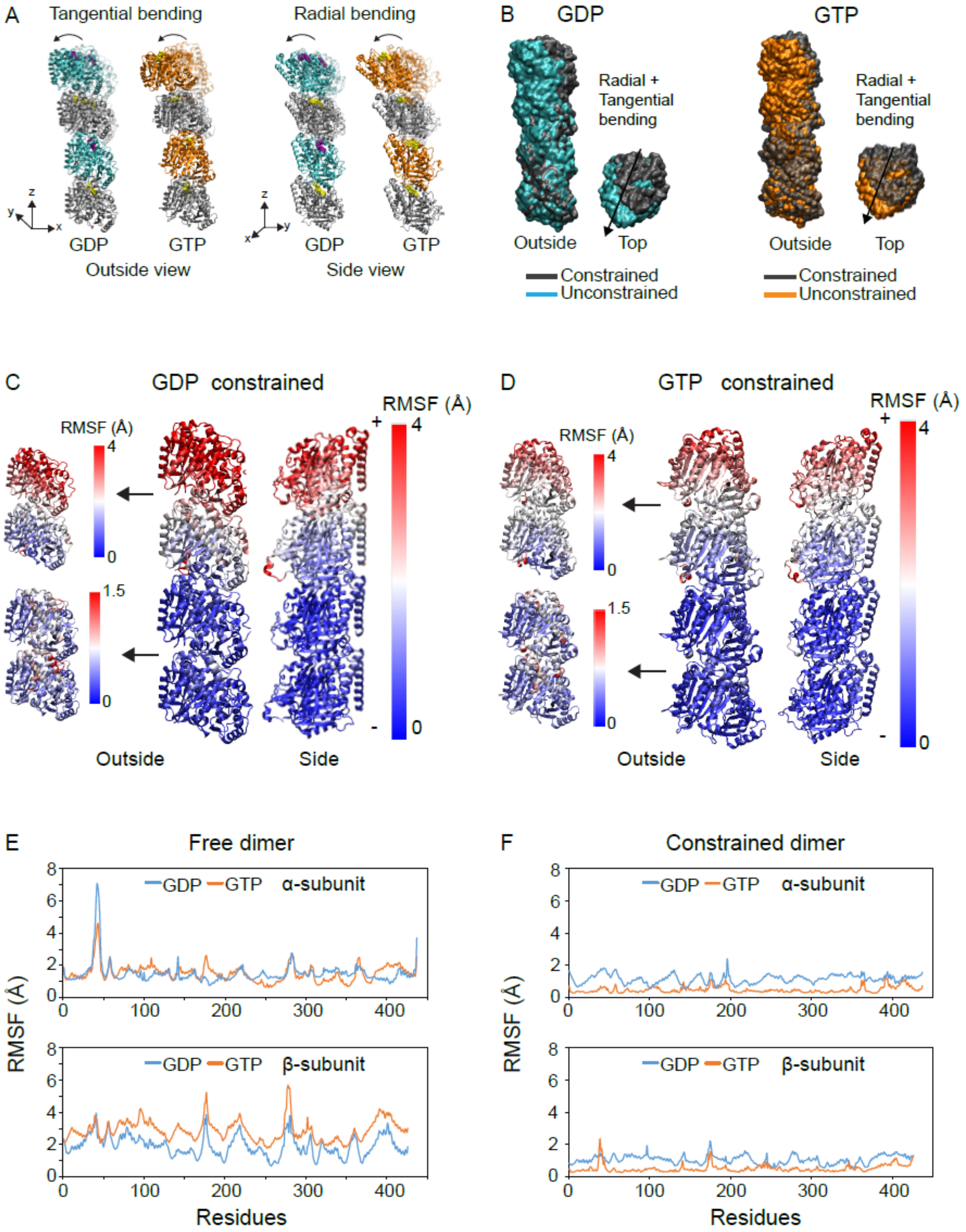


**Figure S1.** Tubulin bending motion analysis indicates two major bending modes: radial and tangential bending in both unconstrained and constrained oligomers. (A) Tubulin oligomer bending directions are indicated with the crystal structure (transparent) and the average structure (opaque). (B) Bending modes of the constrained oligomers are compared to the unconstrained oligomers for both nucleotides. (C) Average RMSF values of constrained tubulin residues are shown from microtubule side and outside views for GDP-tubulin and (D) GTP-tubulin. Color bar shows RMSF values in angstroms. (E) Average RMSF values for all α- and β-subunit residues of the free (top) dimer for GDP- and GTP-tubulin. (F) Average RMSF values for all α- and β-subunit residues of the constrained (bottom) dimer for GDP- and GTP-tubulin. α-tubulin is shown as silver, GDP β-tubulin is shown as cyan and GTP β-tubulin is in orange.

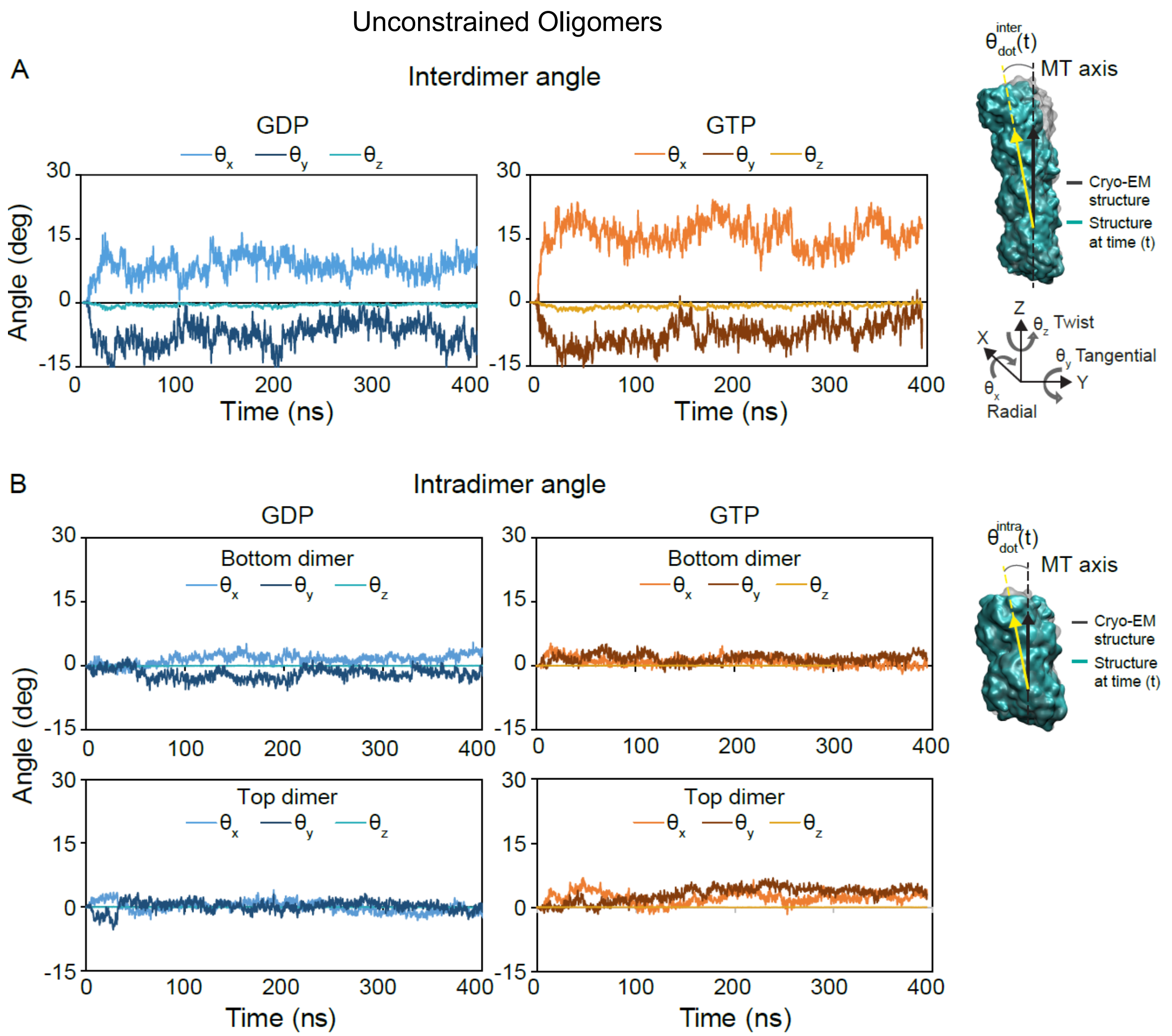


**Figure S2.** Tubulin dimer bending angles for the unconstrained MD trajectories, decomposed into three perpendicular angles, $\theta_x$, radial, $\theta_y$, tangential, and $\theta_z$, twist. (A) interdimer bending angles for GDP- and GTP-tubulin dimers. Right figure indicates how the interdimer angles are calculated after the first dimer's alignment for every time frame. Transparent grey shows the reference structure and cyan shows the structure at time t. (B) Intradimer bending angles for GDP- and GTP-tubulin dimers for the bottom and top dimers. Right figure indicates how the intradimer angles are calculated after the first dimer's alignment for every time frame.

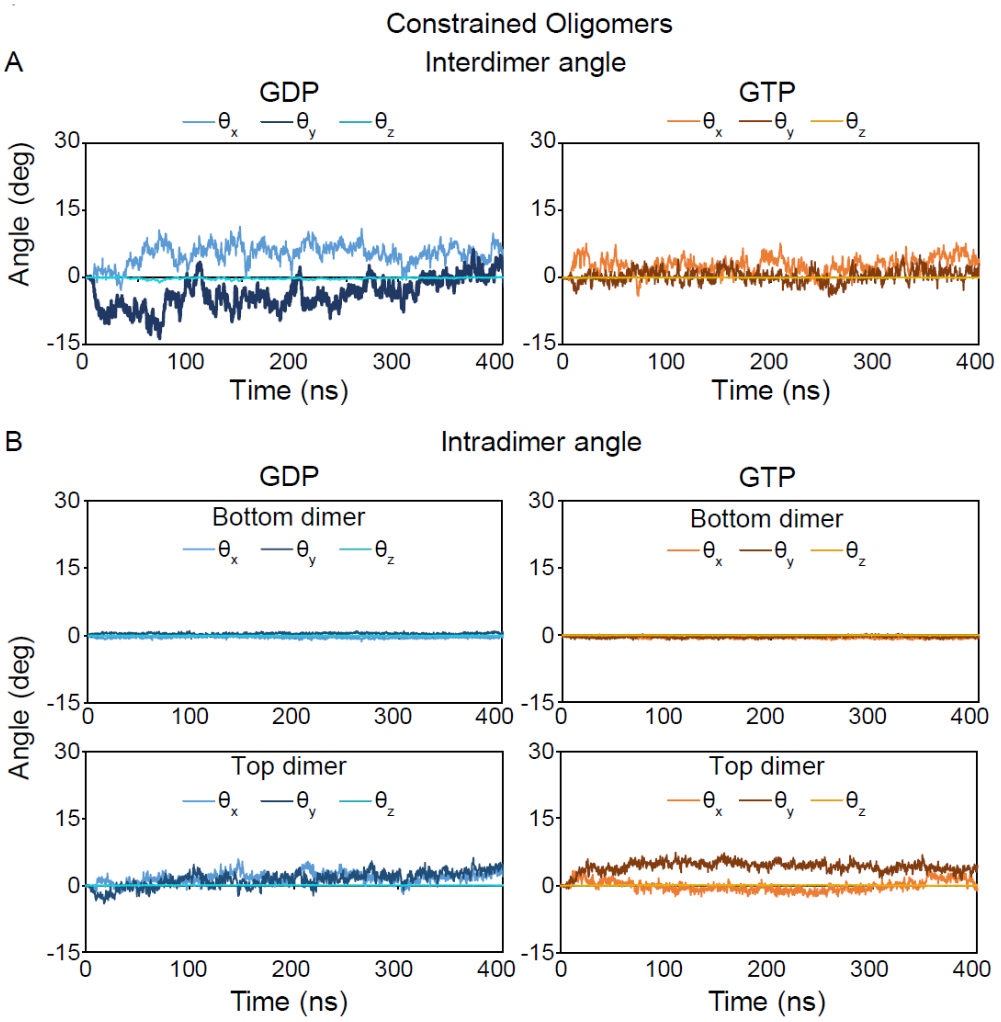


**Figure S3.** Tubulin dimer bending angles shown for the constrained MD trajectories, decomposed into three perpendicular angles, $\theta_x$, radial, $\theta_y$, tangential, and $\theta_z$, twist. (A) interdimer bending angles for GDP- and GTP-tubulin dimers. (B) Intradimer bending angles for GDP- and GTP-tubulin dimers for the bottom (constrained) and top (free) dimers.

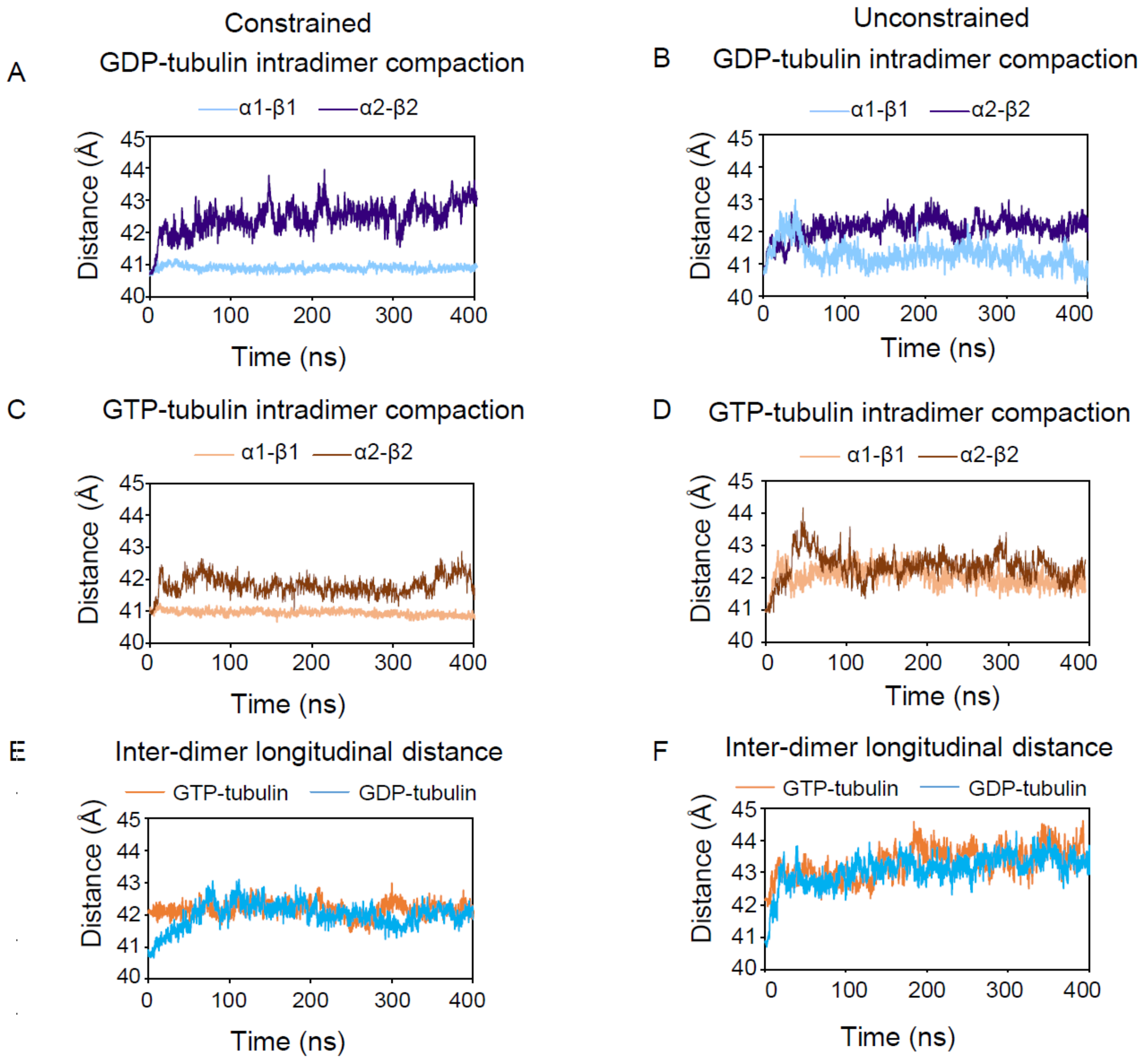


**Figure S4.** Longitudinal compaction release of the dimers is more significant in the already compacted GDP-tubulins. (A, B) intradimer compaction indicated as a function of time for the bottom dimer (α1-β1) and top dimer (α2-β2) for GDP- and (C, D) GTP-tubulin dimers in constrained and unconstrained simulations respectively. (E) Interdimer compaction as a function of time for GDP- and GTP-tubulin dimers in constrained and (F) unconstrained simulations. Compaction is calculated based on COM-to-COM distance of the monomers at different interfaces.

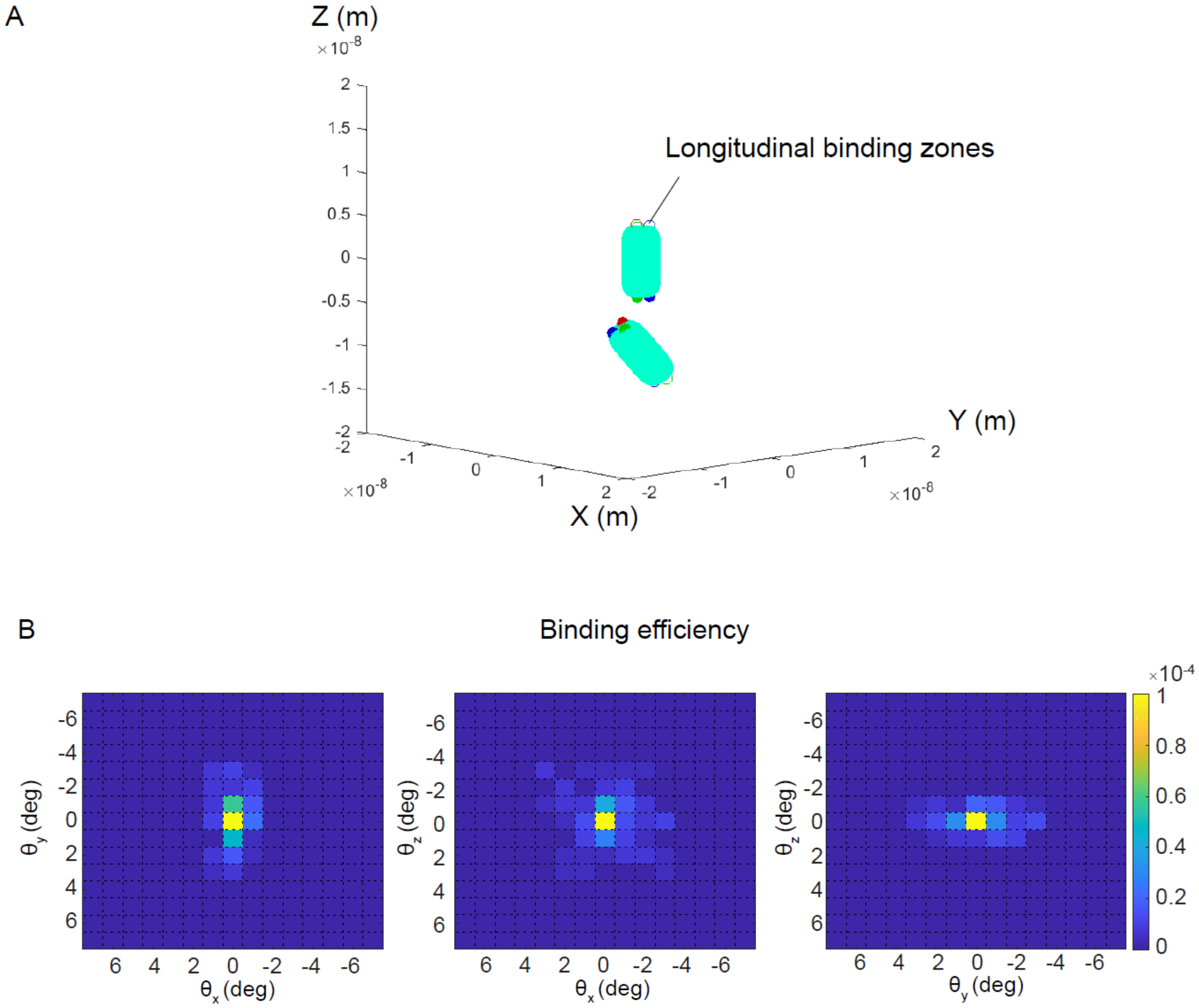


**Figure S5.** BD simulations of two tubulin dimers shows binding efficiency is highest when the dimers approach each other relatively straight on and well-aligned. (A) Representation of the tubulin dimers with three longitudinal binding zones in BD simulations. (B) Binding efficiency as a function of different relative rotational angles of the dimers, with $\theta_x$, radial, $\theta_y$, tangential, and $\theta_z$, twist bending angles. Color bar shows binding efficiency.

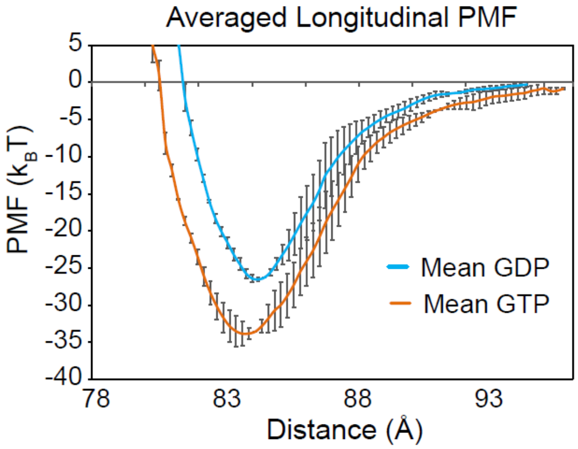


**Figure S6**. Averaged longitudinal PMFs of 15 replicates for GDP- and GTP-tubulin dimers as a function of center-to-center distance demonstrates a stronger bond for GTP dimers. Error bars are standard error of the mean of 15 replicates for each nucleotide.

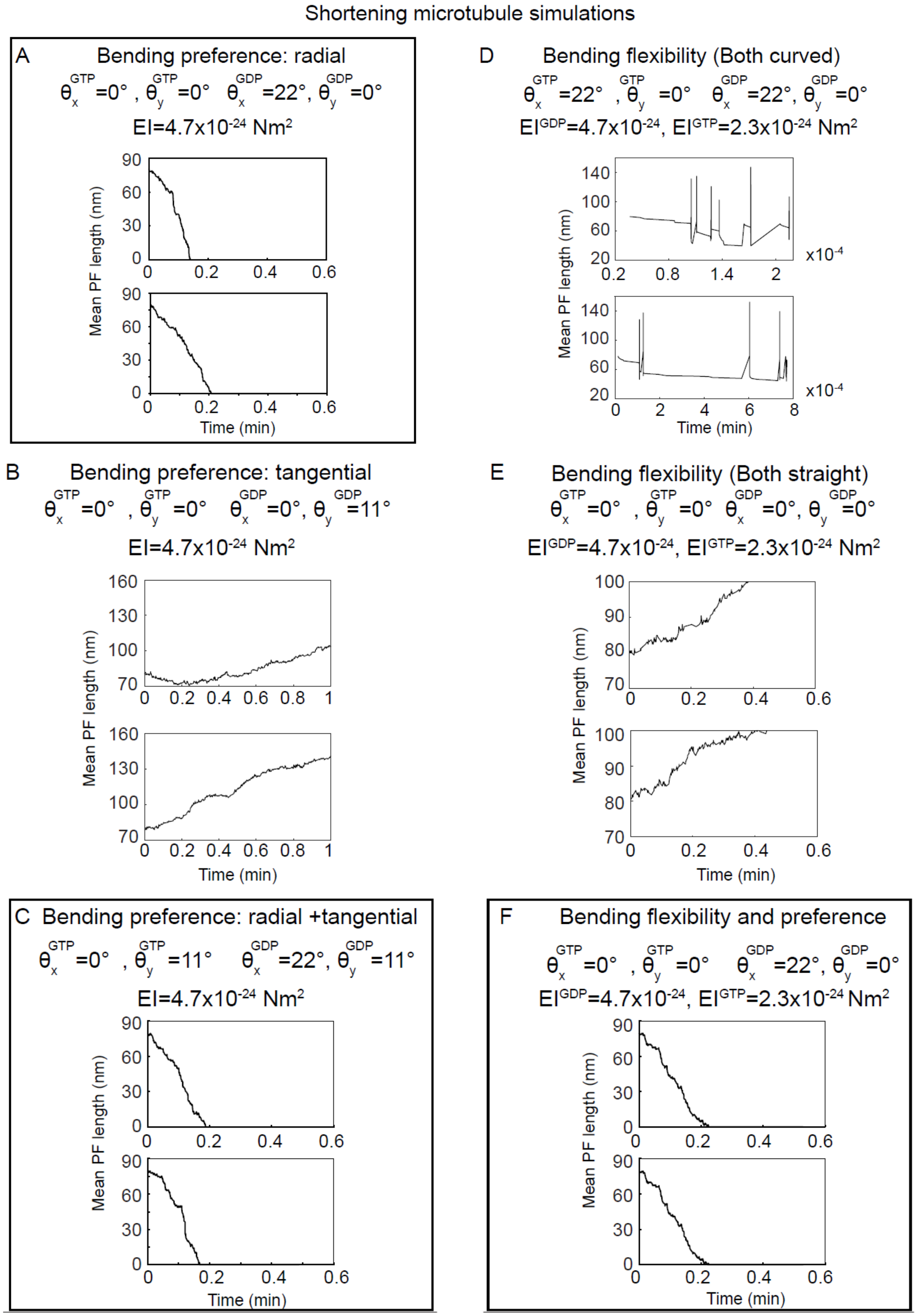


**Figure S7.** Shortening microtubule length-time histories for various conditions of bending preferences and flexibilities indicate shrinkage only when radial bending preference is present. Simulated shortening microtubules started from an uncapped blunt configuration and run for 500 events, with free tubulin concentration of 10 μM. (A) Microtubule length-time is shown when GDP-tubulin has a radially outward bending preference of 22°, GTP-tubulin prefers to be straight, and both dimers have the flexibility of $EI=4.7x10^{-24}$ $Nm^2$. Instant shortening is observed. (B) Microtubule length-time is shown when GDP-tubulin has a tangential bending preference of 11°, GTP-tubulin prefers to be straight, and both dimers have the flexibility of $EI=4.7x10^{-24}$ $Nm^2$. Growing microtubules are mainly observed, and shrinkage happens rarely. (C) Microtubule length-time is shown when GDP-tubulin has a radially outward bending preference of 22°, both GDP- and GTP-tubulin prefers a tangential bending of 11°, and both dimers have the flexibility of $EI=4.7x10^{-24}$ $Nm^2$. Microtubule shortening is observed. (D) Microtubule length-time is shown when GDP-tubulin is stiffer than GTP-tubulin with $EI^{GDP}=4.7x10^{-24}$ $Nm^2$, $EI^{GTP}=2.3$ $x10^{-24}$ $Nm^2$, and both dimers prefer a radially outward bending of 22°. Abnormal dynamics are observed. (E) Microtubule length-time is shown when GDP-tubulin is stiffer than GTP-tubulin with $EI^{GDP}=4.7$ $x10^{-24}$ $Nm^2$ , $EI^{GTP}=2.3$ $x10^{-24}$ $Nm^2$, and both dimers prefer to be straight. Only growing microtubules are observed without shortening events. (F) Microtubule length-time is shown when GDP-tubulin is stiffer than GTP-tubulin with $EI^{GDP}=4.7$ $x10^{-24}$ $Nm^2$ , $EI^{GTP}=2.3$ $x10^{-24}$ $Nm^2$, and GDP dimer has a radial bending preference of 22°. Microtubule shortening is shown. Bold borders show mechanisms consistent with experimental results.

Figure S8 Growing microtubule simulations (13 PFs capped)

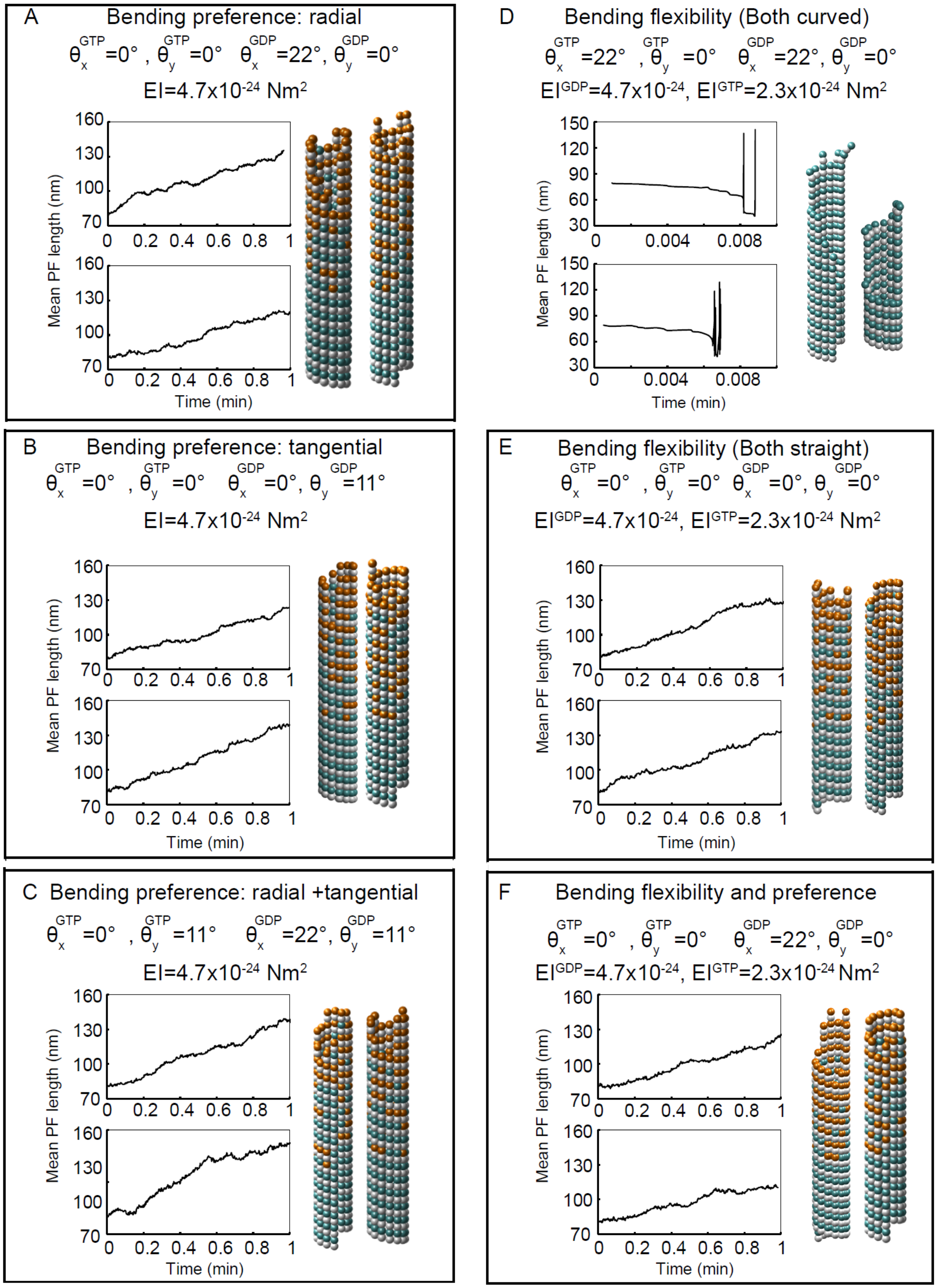

**Figure S8.** Growing microtubule length-time histories for various conditions of bending preferences and flexibilities indicate growing except for when both dimers are kinked with a flexibility difference. Simulated growing microtubules started from a 13-protofilament GTP-capped (4 layers) blunt configuration ([GTP-tubulin]=10 μM) and run for 500 events. Simulated growing microtubule tip structures are shown next to the length-time histories. (A) Microtubule length-time is shown when GDP-tubulin has a radially outward bending preference of 22°, GTP-tubulin prefers to be straight, and both dimers have the flexibility of $EI=4.7x10^{-24}$ $Nm^2$. Microtubule growth is observed. (B) Microtubule length-time is shown when GDP-tubulin has a tangential bending preference of 11°, GTP-tubulin prefers to be straight, and both dimers have the flexibility of $EI=4.7x10^{-24}$ $Nm^2$. Growing microtubule is observed. (C) Microtubule length-time is shown when GDP-tubulin has a radially outward bending preference of 22°, both GDP- and GTP-tubulin prefers a tangential bending of 11°, and both dimers have the flexibility of $EI=4.7x10^{-24}$ $Nm^2$. Growing microtubule is observed. (D) Microtubule length-time is shown when GDP-tubulin is stiffer than GTP-tubulin with $EI^{GDP}=4.7x10^{-24}$ $Nm^2$, $EI^{GTP}=2.3$ $x10^{-24}$ $Nm^2$, and both dimers prefer a radially outward bending of 22°. Abnormal dynamics are observed. (E) Microtubule length-time is shown when GDP-tubulin is stiffer than GTP-tubulin with $EI^{GDP}=4.7$ $x10^{-24}$ $Nm^2$ , $EI^{GTP}=2.3$ $x10^{-24}$ $Nm^2$, and both dimers prefer to be straight. Microtubule growth is observed. (F) Microtubule length-time is shown when GDP-tubulin is stiffer than GTP-tubulin with $EI^{GDP}=4.7$ $x10^{-24}$ $Nm^2$ , $EI^{GTP}=2.3$ $x10^{-24}$ $Nm^2$, and GDP dimer has a radial bending preference of 22°. Growing microtubule is shown. Bold borders show mechanisms consistent with experimental results.

### Supporting Movies

**Movie S1.** Radial bending motion of unconstrained tubulin oligomers in GDP- (left) and GTP-state (right). α-tubulin is depicted in silver and β-tubulin is shown in cyan and orange for GDP- and GTP-states respectively. GTP- and GDP-nucleotide are represented in yellow and purple, respectively. Side view of a microtubule is shown.

**Movie S2.** Tangential bending motion of unconstrained tubulin oligomers in GDP- (left) and GTP-state (right). α-tubulin is depicted in silver and β-tubulin is shown in cyan and orange for GDP- and GTP-states respectively. GTP- and GDP-nucleotide are represented in yellow and purple, respectively. Outside view of a microtubule is shown.
